# A stochastic mathematical model of 4D tumour spheroids with real-time fluorescent cell cycle labelling

**DOI:** 10.1101/2021.11.28.470300

**Authors:** Jonah J. Klowss, Alexander P. Browning, Ryan J. Murphy, Elliot J. Carr, Michael J. Plank, Gency Gunasingh, Nikolas K. Haass, Matthew J. Simpson

## Abstract

*In vitro* tumour spheroid experiments have been used to study avascular tumour growth and drug design for the last 50 years. Unlike simpler two-dimensional cell cultures, tumour spheroids exhibit heterogeneity within the growing population of cells that is thought to be related to spatial and temporal differences in nutrient availability. The recent development of real-time fluorescent cell cycle imaging allows us to identify the position and cell cycle status of individual cells within the growing population, giving rise to the notion of a four-dimensional (4D) tumour spheroid. In this work we develop the first stochastic individual-based model (IBM) of a 4D tumour spheroid and show that IBM simulation data qualitatively and quantitatively compare very well with experimental data from a suite of 4D tumour spheroid experiments performed with a primary human melanoma cell line. The IBM provides quantitative information about nutrient availability within the spheroid, which is important because it is very difficult to measure these data in standard tumour spheroid experiments. Software required to implement the IBM is available on GitHub, https://github.com/ProfMJSimpson/4DFUCCI.

## 1. Introduction

*In vitro* tumour spheroid experiments are widely-adopted to study avascular tumour growth and anti-cancer drug design [1–3]. Unlike simpler two-dimensional assays, tumour spheroid experiments exhibit heterogeneity within the growing population of cells, and this heterogeneity is thought to be partly driven by spatial and temporal differences in the availability of diffusible nutrients, such as oxygen [3, 4]. Historically, tumour spheroids have been analysed experimentally using bright field imaging to measure the size of the growing spheroid [5, 6], however this approach does not reveal information about the internal structure of the growing population. Since 2008, *fluorescent ubiquitination-based cell cycle indicator* (FUCCI) has enabled real-time identification of the cell cycle for individual cells within growing populations [4, 7, 8]. Using FUCCI, nuclei of cells in G1 phase fluoresce red, nuclei of cells in S/G2/M phase fluoresce green, and nuclei of cells in early S (eS) phase appear yellow as a result of both red and green fluorescence being active [7] (Figure 1a). FUCCI simultaneously provides information about spheroid size and heterogeneity of the cell cycle status (Figure 1c-e). In particular, at early times the entire spheroid is composed of freely cycling cells, with a relatively even distribution of FUCCI colours, whereas at intermediate times cells in the central region become predominantly red, indicating G1-arrest [4]. Late time growth is characterised by the formation of a central necrotic region, indicated by a complete absence of fluorescence. FUCCI allows us to identify both the position of individual cells within the growing spheroid in three spatial dimensions, as well as identifying cell cycle status, giving rise to the notion of a *four-dimensional (4D) tumour spheroid* [9]. Assuming spherical symmetry, we can characterise the geometry of 4D spheroids by three radii: *r*_o_(*t*) *>* 0 is the outer radius, *r*_a_(*t*) ≥ 0 is the arrested radius, and *r*_n_(*t*) ≥ 0 is the necrotic radius, with *r*_o_(*t*) *> r*_a_(*t*) ≥ *r*_n_(*t*). In Figure 1e, we see that *r*_n_(*t*) = 0 for *t* ≤ 3, with the necrotic core forming sometime between *t* = 3 and *t* = 6 days.

**Figure 1:**
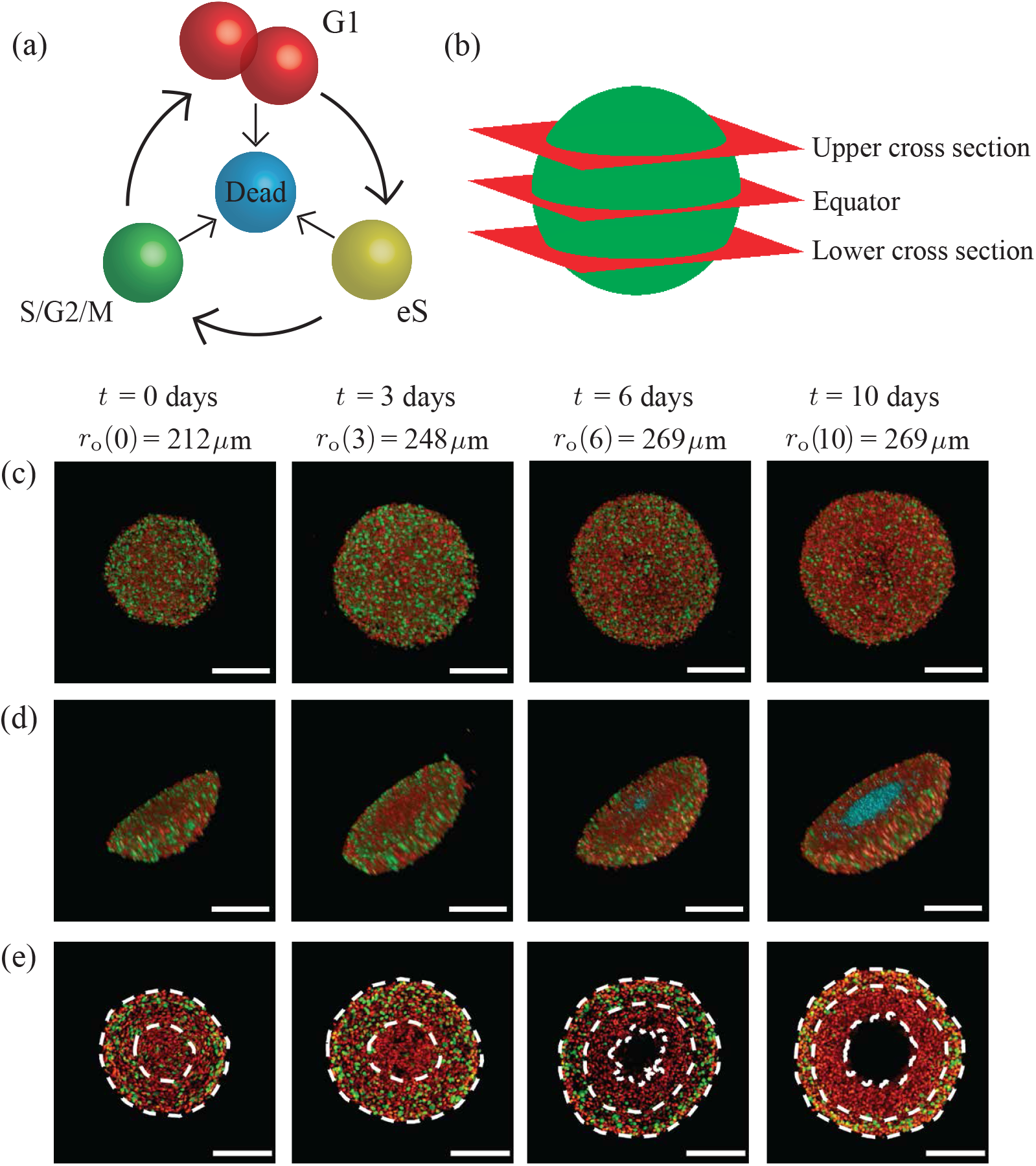
Motivation. (a) A schematic of the cell cycle, indicating the transition between different cell cycle phases, and their associated FUCCI fluorescence. Red, yellow, and green colouring indicates cells in G1, eS, and S/G2/M phase, respectively. (b) Locations of the upper cross section, equator and lower cross section. (c)–(e) Experimental images of a tumour spheroid using the human melanoma cell line WM793B at days 0, 3, 6, and 10 (after formation) showing: (c) full spheroids, viewed from above; (d) spheroid hemispheres; and, (e) spheroid slices, where the cross section is taken at the equator. White dashed lines in (e) denote the boundaries of different regions, where the outermost region is the proliferative zone, the next region inward is the G1-arrested region, and the innermost region at days 6 and 10 is the necrotic core. In (a) and (d) we use cyan colouring for dead cells, which assist in identifying the necrotic core in (d). Spheroid outer radii are labelled alongside their corresponding time points, and scale bars represent 200 *μ*m.

Continuum mathematical models of tumour spheroids have been developed, analysed, and deployed for over 50 years [10–18], and these developments have included very recent adaptations of classical models so that they can be used to study tumour spheroids with FUCCI [9]. However, continuum modelling approaches lack the ability to track individual cells within the growing population, and typically neglect heterogeneity and stochasticity within the population. In comparison, individual-based models (IBMs) allow us to study population dynamics in detail by keeping track of all individuals within the population, as well as explicitly including effects of heterogeneity and stochasticity [19–23]. While some previous IBMs have been developed to describe classical tumour spheroid experiments without FUCCI [24, 25], no IBMs have been developed with the specific goal of simulating 4D tumour spheroid experiments with FUCCI.

In this work, we develop a continuous-space, continuous-time IBM of 4D tumour spheroid growth with FUCCI. The IBM explicitly describes how individual cells migrate, die, and progress through the cell cycle to mimic FUCCI. Certain mechanisms in the IBM are coupled to the local availability of a diffusible nutrient. We demonstrate the biological fidelity of the IBM by qualitatively comparing simulation results with detailed experimental images at several cross sections (Figure 1b), with the aim of providing more comprehensive detail about the internal structure. Quantitative data from the model are then used to assess the spheroid population distribution, nutrient concentration, and the role variability plays in the spheroid. We extract and quantitatively compare simulation radius estimates with measurements from a series of 4D tumour spheroid experiments using a human primary melanoma cell line (Figure 1). Using a careful choice of parameter values, we also show that the IBM quantitatively replicates key features of 4D tumour spheroids.

## 2. Methods

### 2.1. Experimental methods

#### Spheroid growth and staining

Human melanoma cells from the WM793B cell line were genotypically characterised [26–28], grown as described in [3], and authenticated by short tandem repeat fingerprinting (QIMR Berghofer Medical Research Institute, Herston, Australia). The WM793B cells were transduced with FUCCI constructs [4]. Spheroid seeding, growth, and staining were performed as described in [3], with 1% penicillin-streptomycin (ThermoFisher, Massachusetts, USA). Three 96-well plates of spheroids, seeded with a density of 10,000 cells per well, were grown and harvested over 14 days. One 96-well plate was placed in an IncuCyte S3 (Sartorius, Göttingen, Germany) and imaged at 6 hour intervals over 14 days. Harvested spheroids were stained with either DRAQ7 (ThermoFisher, Massachusetts, USA) for necrosis or pimonidazole for hypoxia, fixed in 4% paraformaldehyde solution, and stained with DAPI as per [29].

To reveal the hypoxic region, spheroids stained with pimonidazole were permeabilised with 0.5% triton X-100 in phosphate buffered solution (PBS) for one hour, then blocked in antibody dilution buffer (Abdil) [30] for 24 hours. Spheroids were stained with a 1:50 anti-pimonidazole mouse IgG1 monoclonal antibody (Hypoxyprobe-1 MAb1) in Abdil for 48 hours, before washing in PBS with 0.1% tween-20 for six hours. These spheroids were then placed in a 1:100 solution of Alexa Fluor 647 in Abdil for 48 hours. Following this, the spheroids were washed for six hours in PBS.

#### Confocal imaging

Harvested spheroids were mounted in 2% low melting agarose in PBS solution and cleared in clearing reagent 2 with matching refractive index [29], on #1.5 glass bottom plates. For collecting 2D cross sections, images were taken at the equator and upper and lower cross sections (Figure 1b), which we define as the Z coordinate halfway between the equator and the top or bottom of the spheroid. If the necrotic core exists, the upper and lower cross sections are at the top or bottom of the necrotic core, respectively. 3D spheroid images were collected by imaging over the entire Z range of the spheroid.

#### Computational image analysis

The image processing algorithm [31] was used to estimate *r*_o_(*t*), *r*_a_(*t*), and *r*_n_(*t*).

### 2.2. Individual-based mathematical model

We simulate 4D spheroid growth inside a cubic domain, Ω, of side length *L*, where *L* is chosen to be large enough so that agents do not reach the boundary of the domain during the simulation, but not so large as to incur significant computational overhead (Supplementary S3.3). Biological cells are represented as discrete agents located at **x**_*n*_(*t*) = (*x*_*n*_(*t*), *y*_*n*_(*t*), *z*_*n*_(*t*)) for *n* = 1, 2, 3, …, *N* (*t*), where *N* (*t*) is the total number of agents at time *t*.

#### Gillespie algorithm

The IBM describes key cellular-level behaviours; namely cell cycle progression and mitosis, cell motility, and cell death, as discrete events simulated using the Gillespie algorithm [32]. Each agent has an allocated rate of cell cycle progression, dependent on its cell cycle status and the local nutrient concentration (Figure 2a). Agents in each phase of the cell cycle are coloured according to FUCCI, with G1 agents coloured red, eS agents coloured yellow, and S/G2/M agents coloured green.

**Figure 2:**
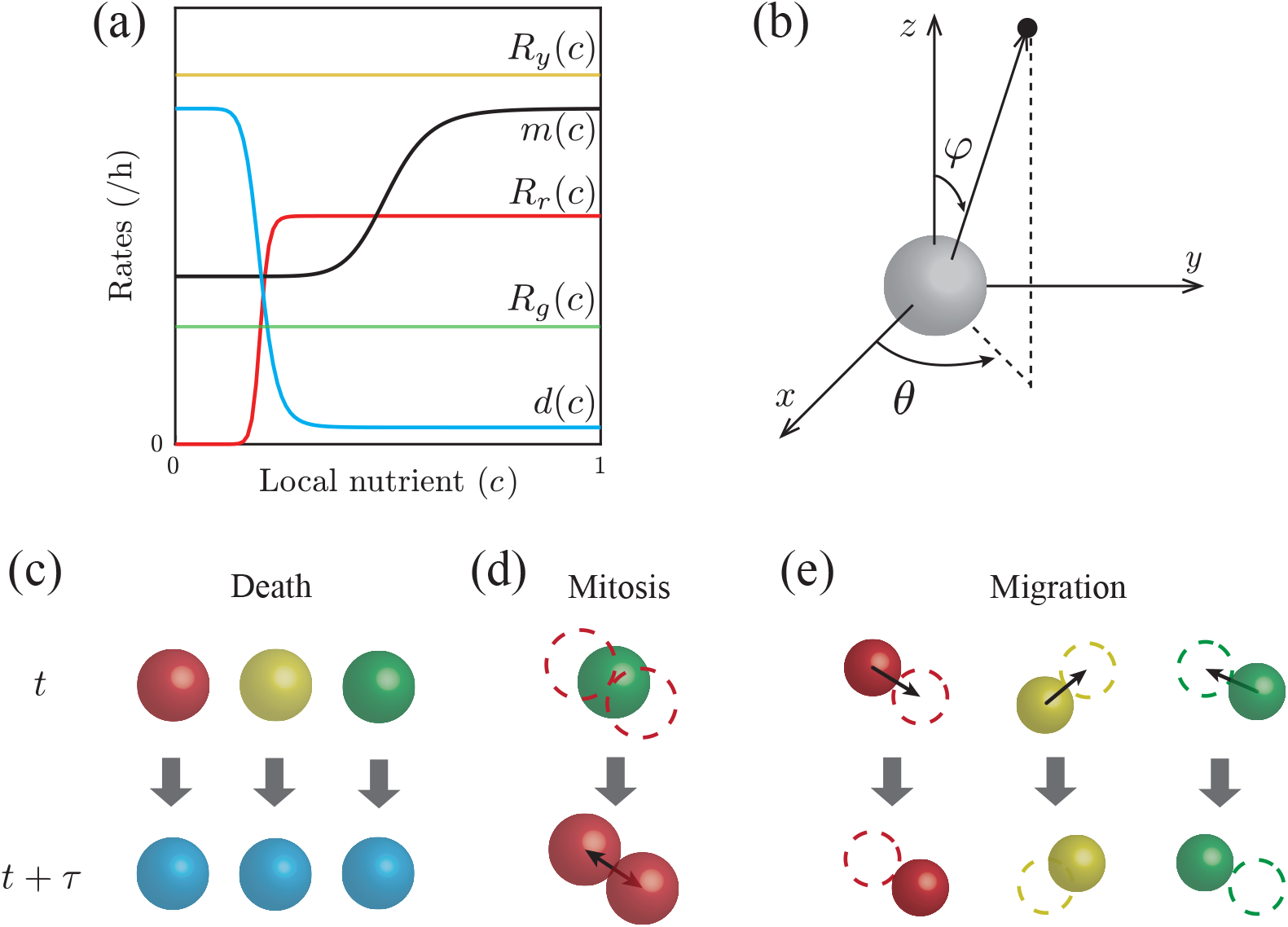
IBM schematic. (a) Nutrient-dependent rates (Equations (1)–(5)). (b) Random directions for migration and mitosis are obtained by sampling the polar angle *θ*, and the azimuthal angle *ϕ* separately [33]. (c)–(e) Schematics showing agent-level events; death, mitosis, and migration, across a time interval of duration *τ*. (c) Any living agent may die, removing it from the simulation. (d) An agent located at **x**_*n*_ undergoes mitosis to produce two daughter agents in G1 phase and dispersed a distance of *σ/*2 from **x**_*n*_ in opposite, randomly chosen directions. (e) Any living agent can migrate in a random direction with step length *μ*.

We make the natural assumption that biological cells require access to sufficient nutrients to commit to entering the cell cycle. Therefore, the red-to-yellow transition rate, *R*_*r*_(*c*), depends on the local nutrient concentration, *c*(**x**, *t*) (Figure 2a). Once an agent has committed to entering the cell cycle, we assume the yellow-to-green transition takes place at a constant rate *R*_*y*_, and the green-to-red transition, which involves mitosis, occurs at a constant rate *R*_*g*_ (Figure 2a).

The rate of agent death is assumed to depend on the local nutrient concentration, *d*(*c*). When an agent dies, it is removed from the simulation and we record the location at which the death event occurs (Figure 2c). When an agent moves or undergoes mitosis (Figure 2d-e), a random direction in which the agent will migrate, or its daughter agents will disperse, is chosen (Figure 2b). For an agent undergoing mitosis, the first daughter agent is placed a distance *σ/*2 along the randomly chosen direction, and the second daughter agent is placed at a distance *σ/*2 in the opposite direction, leaving the two daughter agents dispersed a distance of *σ* apart, where we set *σ* to be equal to a typical cell diameter [34] (Figure 2d, Table 1). When migrating, agents are displaced a distance *μ*along the randomly chosen direction (Figure 2e). Similar to the dispersal, we simulate migration by taking the step length *μ*to be a typical cell diameter.

**Table 1:** IBM parameter values.

| Parameter Name | Symbol | Value | Source |
| --- | --- | --- | --- |
| *Numerical Parameters* |  |  |  |
| Initial number of agents | $N(0)$ | 30 000 | Experimental measurement<br>(Supplementary S6) |
| Initial number of red agents | $N_r(0)$ | 20,911 | Assumption<br>(Supplementary S7) |
| Initial number of yellow agents | $N_y(0)$ | 995 | Assumption<br>(Supplementary S7) |
| Initial number of green agents | $N_g(0)$ | 8,094 | Assumption<br>(Supplementary S7) |
| Domain length | $L$ | 4000 $\mu\text{m}$ | Numerical experiments<br>(Supplementary S3.3) |
| Initial spheroid radius | $r_o(0)$ | 245 $\mu\text{m}$ | Experimental measurement |
| Dispersal distance | $\sigma$ | 12 $\mu\text{m}$ | Assumption<br>(Supplementary S4) |
| Migration distance | $\mu$ | 12 $\mu\text{m}$ | Assumption<br>(Supplementary S4) |
| Simulation termination time | $T$ | 240 h | Experimental measurement |
| *Per Capita Agent Rates* |  |  |  |
| Maximum G1-eS transition rate | $R_r$ | 0.047 /h | Experimental measurement<br>(Supplementary S5) |
| Constant eS-S/G2/M transition rate | $R_y$ | 0.50 /h | Experimental measurement<br>(Supplementary S5) |
| Constant S/G2/M-G1 transition rate (mitosis) | $R_g$ | 0.062 /h | Experimental measurement<br>(Supplementary S5) |
| Maximum death rate | $d_{\max}$ | 2 /h | Assumption |
| Minimum death rate | $d_{\min}$ | 0.0005 /h | Assumption |
| Maximum migration rate | $m_{\max}$ | 0.12 /h | Assumption |
| Minimum migration rate | $m_{\min}$ | 0.06 /h | Assumption |
| Hill function index for arrest | $\eta_1$ | 5 | Assumption |
| Hill function index for migration | $\eta_2$ | 5 | Assumption |
| Hill function index for death | $\eta_3$ | 15 | Assumption |
| *Nutrient Parameters* |  |  |  |
| Number of nodes | $I^3$ | 201<sup>3</sup> | Assumption<br>(Supplementary S3.4) |
| Steady-state solution interval | $t^*$ | 1 h | Assumption<br>(Supplementary S3.4) |
| Consumption-diffusion ratio | $\alpha$ | 0.15<br>$\mu\text{m}/\text{cells}$ | Assumption |
| Critical arrest concentration | $c_a$ | 0.4 | Assumption |
| Critical migration concentration | $c_m$ | 0.5 | Assumption |
| Critical death concentration | $c_d$ | 0.1 | Assumption |

We specify the agent cycle progression rates,

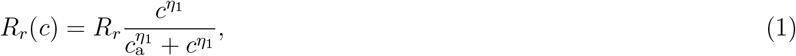

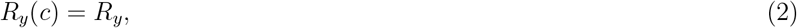

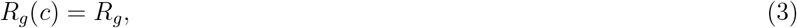

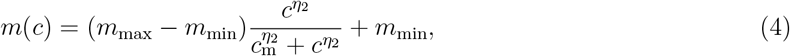

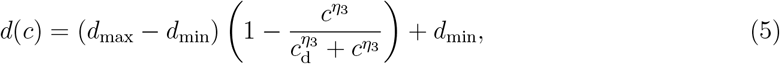

where *c*(**x**_*n*_, *t*) ∈ [0, 1] is the non-dimensional nutrient concentration at the location of the *n*th agent; *R*_*r*_ *>* 0 is the the maximum red-to-yellow transition rate; *m*_max_ *> m*_min_ ≥ 0 are the maximum and minimum migration rates, respectively; *d*_max_ *> d*_min_ ≥ 0 are the maximum and minimum death rates, respectively; *η*_1_ *>* 0, *η*_2_ *>* 0, and *η*_3_ *>* 0 are Hill function indices; and *c*_a_ *>* 0, *c*_m_ *>* 0, and *c*_d_ *>* 0 are the inflection points of *R*_*r*_(*c*), *m*(*c*), and *d*(*c*) respectively (Figure 2a).

#### Nutrient dynamics

We make the simplifying assumption that cell migration, death, and progression through the cell cycle are regulated by a single diffusible nutrient, such as oxygen [4, 10, 12]. The spatial and temporal distribution of nutrient concentration, *C*(**x**, *t*), is assumed to be governed by a reaction-diffusion equation

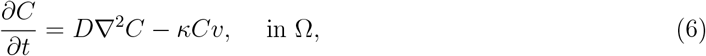

with diffusivity *D >* 0 [*μ*m^2^*/*h], and consumption rate *κ >* 0 [*μ*m^3^*/*(h cells)], and where *v*(**x**, *t*) ≥ 0 [cells*/μ*m^3^] is the density of agents at position **x** and time *t*. The source term in Equation (6) describes the consumption of nutrient at a rate of *κ* [*μ*m^3^*/*(h cells)]. To solve this reaction-diffusion equation we set *v*(**x**_*i,j,k*_, *t*) = *N*_*i,j,k*_*/h*^3^, where *N*_*i,j,k*_ is the number of agents within the control volume surrounding the node located at (*x*_*i*_, *y*_*j*_, *z*_*k*_) and *h*^3^ is the volume of the control volume. On the boundary, ∂Ω, we impose *C* = *C*_b_, where *C*_b_ is some maximum far-field concentration.

Our experiments lead to spheroids of diameter 500–600 *μ*m over a period of 10 days after spheroid formation (Figure 1) (14 days after seeding). Since these length and time scales are clear, we leave the independent variables **x** and *t* in Equation (6) as dimensional quantities. In contrast, spatial and temporal variations of *C*(**x**, *t*) are very difficult to measure during spheroid growth, so we nondimensionalise the independent variable *c*(**x**, *t*) = *C*(**x**, *t*)*/C*_b_, giving

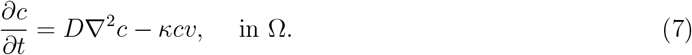

with *c* = 1 on ∂Ω, and *c*(**x**, *t*) ∈ [0, 1].

Typically, the time scale of nutrient diffusion is much faster than the time scale of spheroid growth [10]. Consequently, we approximate Equation (7) by

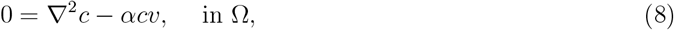

where *α* = *κ/D >* 0 [*μ*m*/*cells]. Therefore, we describe the spatial and temporal distribution of nutrients by solving Equation (8) repeatedly during the simulation. This quasi-steady approximation is computationally convenient, as we describe later. We solve Equation (8) with a finite volume method on a uniform structured mesh (Supplementary S3).

### 2.3. Simulation algorithm

We simulate spheroid growth by supposing the spheroid initially contains *N*(0) agents distributed uniformly within a sphere of radius *r*_o_(0) *>* 0 [*μ*m]. While it is experimentally relevant to assume the population is spherically symmetric at *t* = 0, this assumption is not necessary, and we will discuss this point later. The proportion of agents chosen to be red, yellow, or green at *t* = 0 can be selected arbitrarily, but we choose these proportions so that the internal structure and composition of the *in silico* spheroids are consistent with our *in vitro* measurements. We achieve this by choosing the initial red, yellow, and green population, *N*_r_(0), *N*_y_(0), and *N*_g_(0), respectively, noting that *N*(0) = *N*_r_(0) + *N*_y_(0) + *N*_g_(0) (Supplementary S7). The most appropriate time scale for individual cell-level behaviour is hours, however spheroid development takes place over 10 days, so we will use a mixture of time scales to describe different features of the experiments and simulations as appropriate. We simulate spheroid growth from *t* = 0 to *t* = *T* h, updating the nutrient concentration at *M* equally-spaced points in time. This means that the nutrient concentration is updated at intervals of duration *t*^*∗*^ = *T/M* [h]. The accuracy of our algorithm increases by choosing larger *M* (smaller *t*^*∗*^), but larger *M* decreases the computational efficiency. We explore this tradeoff and find that setting *t*^*∗*^ = 1 h is appropriate (Supplementary S3.4). When Equation (8) is solved for *c*(**x**, *t*), the value of *c*(**x**_*n*_, *t*) at each agent is calculated using linear interpolation. These local nutrient concentrations are held constant for each agent while resolving all the various agent-level events (cycling and proliferation, migration, death) from time *t* to time *t*+*t*^*∗*^. After resolving the appropriate agent-level events, we update the agent density before updating the nutrient profile again. Pseudo-algorithms for the IBM are provided (Supplementary S8), and code to reproduce key results is available on GitHub.

### 2.4. IBM image processing

To estimate *r*_o_(*t*), *r*_a_(*t*), and *r*_n_(*t*), we apply methods described in [18, 31, 35] to the IBM output. Briefly, we import the agent locations from a particular cross section, and map these locations to an (*L* + 1) × (*L* + 1) pixel image, increase the size of the agents to 12 pixels in diameter, and use edge detection to identify and estimate *r*_o_(*t*), *r*_a_(*t*), and *r*_n_(*t*) (Supplementary S1). This procedure adapts the image processing approach for the experimental images so that it is applicable to the synthetic results from the IBM.

## 3. Results and Discussion

We now compare and analyse images and measurements from a range of *in vitro* experiments and *in silico* simulations. All experiments use the WM793B melanoma cell line, which takes approximately four days to form spheroids after the initial seeding in the experiments [36]. This means that *t* = 0 days corresponds to four days after seeding to give the experimental spheroids sufficient time to form. Snapshots from the IBM correspond to a single realisation, however time-series data from the IBM are reported by simulating 10 realisations of the IBM and then averaging appropriate measurements across the 10 simulations.

### 3.1. Parameter values

Table 1 summarises the parameter values used in this study. While some parameters are based on separate, independent two-dimensional experimental measurements (Supplementary S4 – S5) or measurements directly from the spheroids where possible (Supplementary S6), other parameters are chosen based on a series of numerical screening tests (Supplementary S3). We will return to discuss other options for parameter choices later.

### 3.2. Qualitative comparison of experiments and simulations

We now qualitatively compare images of *in vitro* (Figure 3a,c,e) and *in silico* (Figure 3b,d,f) spheroids by imaging various cross sections at different locations, including the equator (Figure 3a-b), the lower cross section (Figure 3c-d), and the upper cross section (Figure 3e-f). We use the definitions in Section 2.1 (Confocal imaging) to identify the lower and upper cross sections in the analysis of both the experimental images and the simulation images. While previous studies have often compared model predictions with experimental observations at a single cross section [25, 36], we aim to provide more comprehensive information about the internal structure of the spheroid by making comparisons at multiple locations.

**Figure 3:**
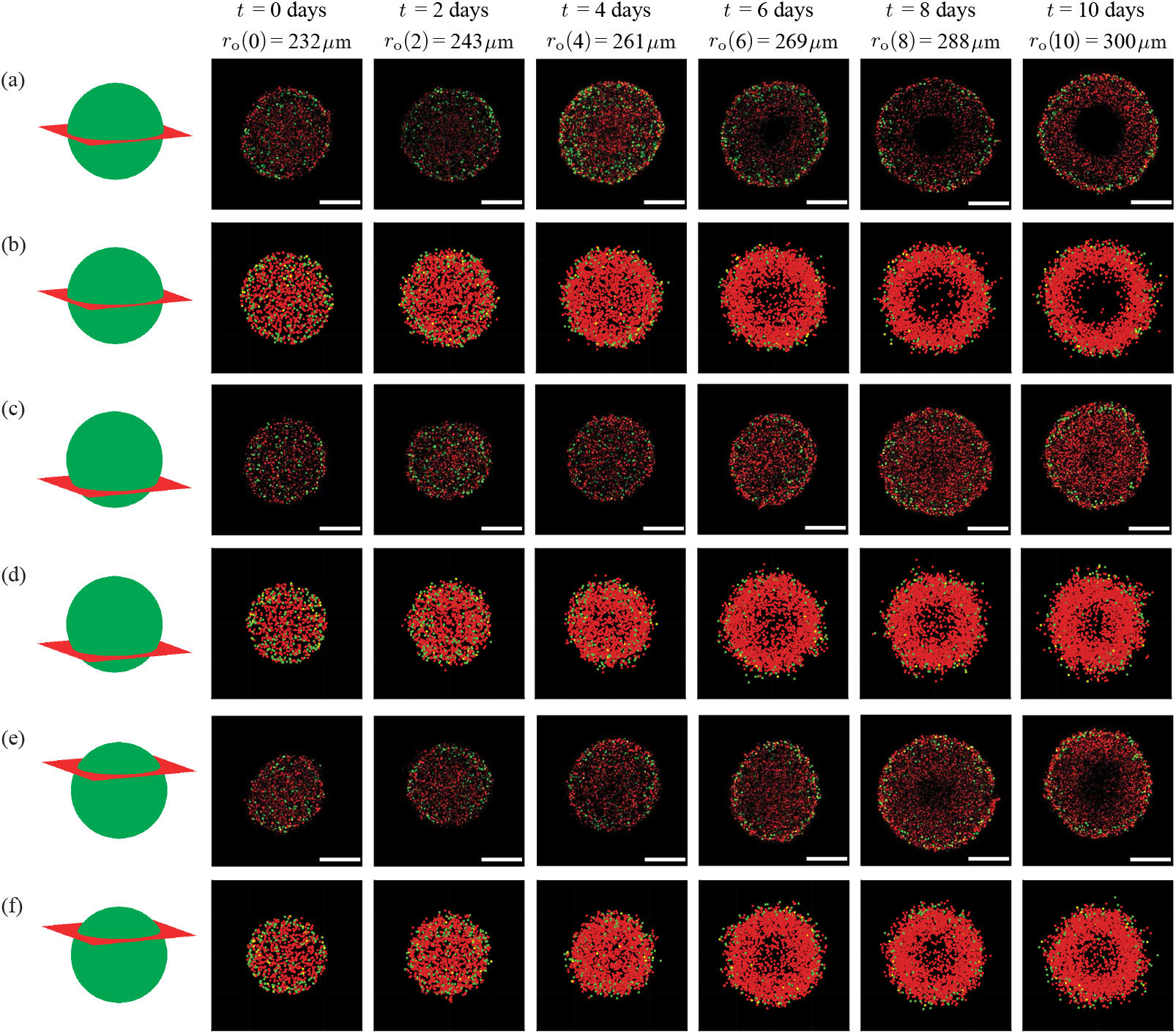
Comparison of *in vitro* and *in silico* 4D spheroids. Experimental results (a,c,e) are compared with simulation results (b,d,f) by examining 2D slices at the equator, lower and upper cross section, respectively. Agent colour (red, yellow, green) corresponds to FUCCI labelling (G1, eS, S/G2/M). Schematics in the left-most column indicate the location of the 2D cross section. The images are taken at (a)–(b) the equator, (c)–(d) the lower cross section, and (e)–(f) the upper cross section. Experimental spheroid radii at the equator are labelled at each time point, and scale bars represent 200 *μ*m.

At the beginning of the experiment, in all cross sections (*in vitro* and *in silico*) we see the population is relatively uniform, with an even distribution of colours, suggesting the entire spheroid is composed of freely-cycling cells. At *t* = 2 and *t* = 4 days, however, we begin to see the development of heterogeneity within the growing *in vitro* and *in silico* populations, with those cells and agents at the centre of the growing spheroid predominantly red, indicating G1-arrest. By *t* = 4 days we see the value of comparing different cross sections, since the G1-arrest is clear in the centre of the equatorial cross section, but there is no obvious heterogeneity present across either the upper or lower cross section at that time. Similarly, by *t* = 6 days we see the formation of a necrotic core in the equatorial cross section, but this is not present at either cross section. By *t* = 8 and *t* = 10 days the spheroid has developed into a relatively complicated heterogeneous structure where the outer spherical shell contains freely cycling cells, the intermediate spherical shell contains living G1-arrested cells, and the internal region does not contain any fluorescent cells.

Overall, the qualitative match between the IBM and the experiment confirms that the IBM captures both the macroscopic growth of the entire spheroid, as well as the emergent spatial and temporal heterogeneity. We now build on this preliminary qualitative information by extracting quantitative measurements of the spheroid growth and exploring the performance of the IBM.

### 3.3. Spheroid structure and nutrient profiles

Given the ability of the IBM to capture key spatial and temporal patterns of spheroid growth, cell cycle arrest, and cell death throughout the spheroid, we now demonstrate how to take these preliminary simulations and extract detailed quantitative data that would be difficult to obtain experimentally. Figure 4a shows a typical IBM simulation during the interval where we observe the development of internal structure. For clarity, we plot the locations of all living agents as in Figure 3, but we now also plot the locations at which agents die, which is difficult to estimate experimentally, but is straightforward with the IBM. Each spheroid in Figure 4a is shown with an octant removed to highlight the development of the internal structure, and for further clarity we show equatorial cross sections in Figure 4b.

**Figure 4:**
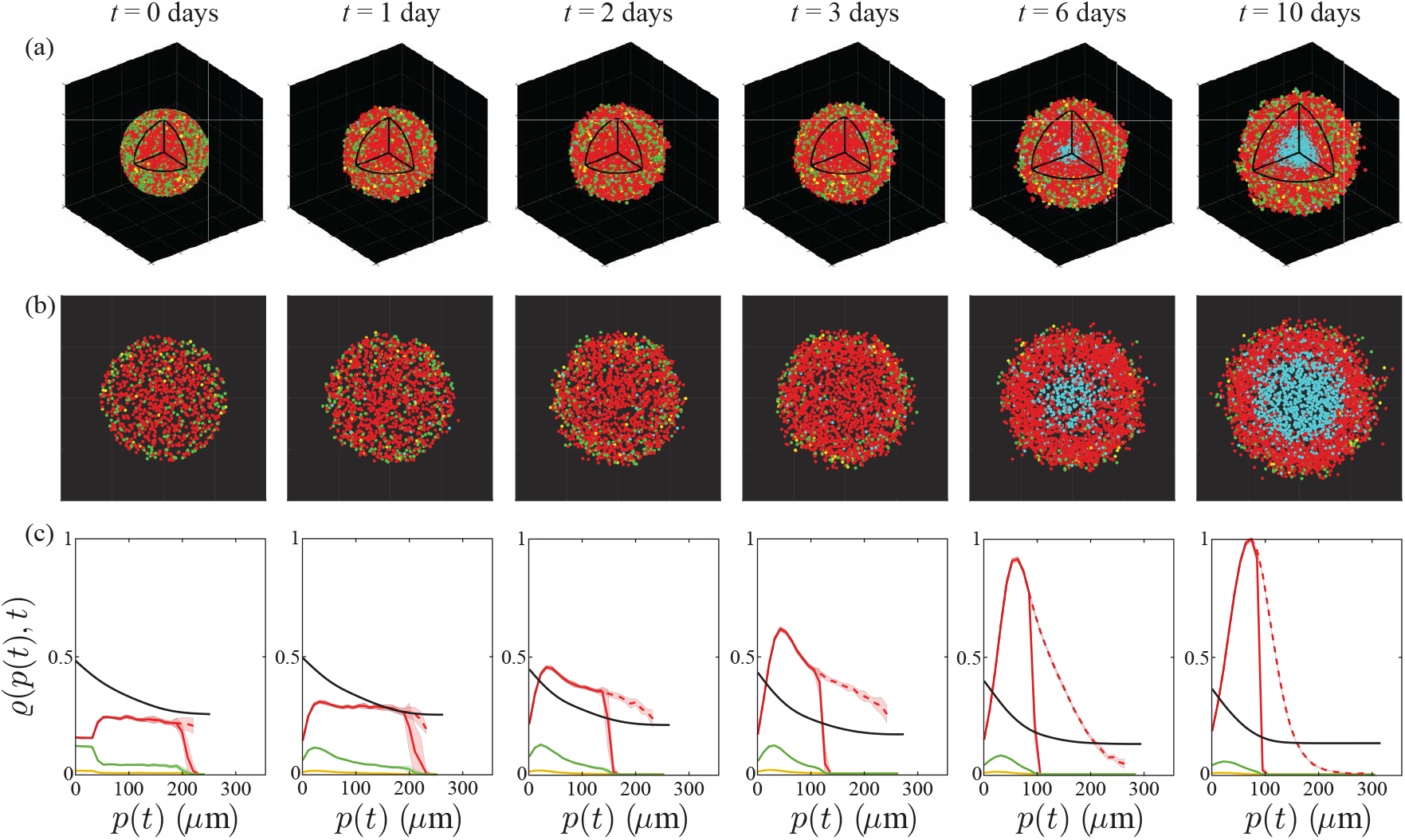
Typical IBM simulation, showing: (a) visualisations of *in silico* spheroids including dead agents (cyan) and (b) cross sections through the spheroid equator with dead agents. (c) Relative concentrations *ϱ*(*p, t*) of nutrient (black) and cycling red, yellow, and green agents (coloured appropriately), based on distance from the periphery *p*(*t*) = *r*_o_(*t*) − *r*, averaged over 10 identically-prepared simulations. The dashed red line shows the relative density of arrested red agents, also averaged over 10 simulations with identical initial conditions. For nutrient, *ϱ*(*p, t*) = *c*. For agents, *ϱ*(*p, t*) is the relative agent density (Supplementary S9). Shaded areas represent plus or minus one standard deviation about the mean, and are non-zero as a consequence of stochasticity in the model, even though the 10 simulations start with identical populations and radii.

To quantify the internal spheroid structure we simulate 10 identically prepared realisations of the IBM and extract averaged quantitative data that are summarised in Figure 4c (Supplementary S9). These data include plotting the nondimensional nutrient concentration, *c*(**x**, *t*), and various normalised agent densities, *ϱ*(*p*(*t*), *t*), as a function of distance from the spheroid periphery, *p*(*t*) = *r*_o_(*t*) − *r*, where *r* is the distance from the spheroid centre. Hence, *p*(*t*) = 0 at the spheroid periphery, and *p*(*t*) = *r*_o_(*t*) at the spheroid centre. This representation of internal spheroid structure is made by assuming that the growing population remains spherically symmetric, which is a reasonable assumption since our initial condition and spheroid growth is spherically symmetric (Figure 4a). Each density profile is normalised relative to the maximum value of all agent densities across all time points, so that we can compare how the density of the various subpopulations of agents and nutrient are distributed (Supplementary S9). Using the IBM we are able to describe the spatial and temporal densities of living agents in various phases of the cell cycle (G1, eS and S/G2/M) as well as G1-arrested agents. We plot each density profile as a function of the distance from the periphery as this allows us to compare various profiles as the size of the spheroid increases [9, 37].

Averaged relative agent density profiles from the IBM provide quantitative information that cannot be easily obtained from experimental observations. Initially we see the relatively evenly distributed G1, eS and S/G2/M populations become rapidly dominated by agents in G1 phase, which then form an obvious inner-most arrested region by about *t* = 2 days. During the interval 3 *< t <* 6 days we see rapid growth in the arrested population, and the eventual formation of a clear necrotic core in the interval 6 *< t <* 10 days. These results indicate the spatial and temporal role of stochasticity, with the variability most evident in the G1 and arrested G1 populations at early times. Plotting the relative agent densities in this way provides a simple approach to interpret the spatial and temporal organisation of cell cycle status within the growing spheroid, and visualising the agent densities together with the nondimensional nutrient concentration is particularly useful when this kind of information cannot be easily obtained experimentally. In particular, it is technically challenging to measure absolute concentrations of nutrient profiles during these experiments [17, 38, 39] and so we now focus on visualising the nutrient concentration profile that drives this heterogeneity. Results in Figure 5 show spatial and temporal patterns in the nutrient profile, *c*(**x**, *t*), for a typical IBM simulation from Figure 4. Figure 5a shows the three-dimensional evolution of *c*(**x**, *t*), with the colourbar highlighting the death and arrest thresholds, *c*_d_ and *c*_a_, respectively. These threedimensional plots show the depletion of nutrient over time in the central region of the spheroid, leading to strong spatial gradients of nutrient concentration near the edge of the growing spheroid. Profiles in Figure 5b show the nutrient profile at the equatorial plane with the *c*(*x, y*, 0) = *c*_a_ contour (red) and the approximate size of the necrotic core (cyan) superimposed. Simplified one-dimensional profiles of *c*(**x**, *t*), along **x** = (*x*, 0,0), are shown in Figure 5c, where the diameter of the growing spheroid (−*r*_o_(*t*) *< x < r*_o_(*t*)) is shaded in yellow. Again, these simplified cross sections illustrate how nutrient consumption leads to the formation of spatial nutrient gradients near the outer radius of the growing spheroid. Overall, a key strength of the IBM is the ability to extract agent-level information (Figure 4) as well as information about the nutrient distribution (Figure 5), whereas experimental studies typically report cell-level data without explicitly showing nutrient-level information [4, 6].

**Figure 5:**
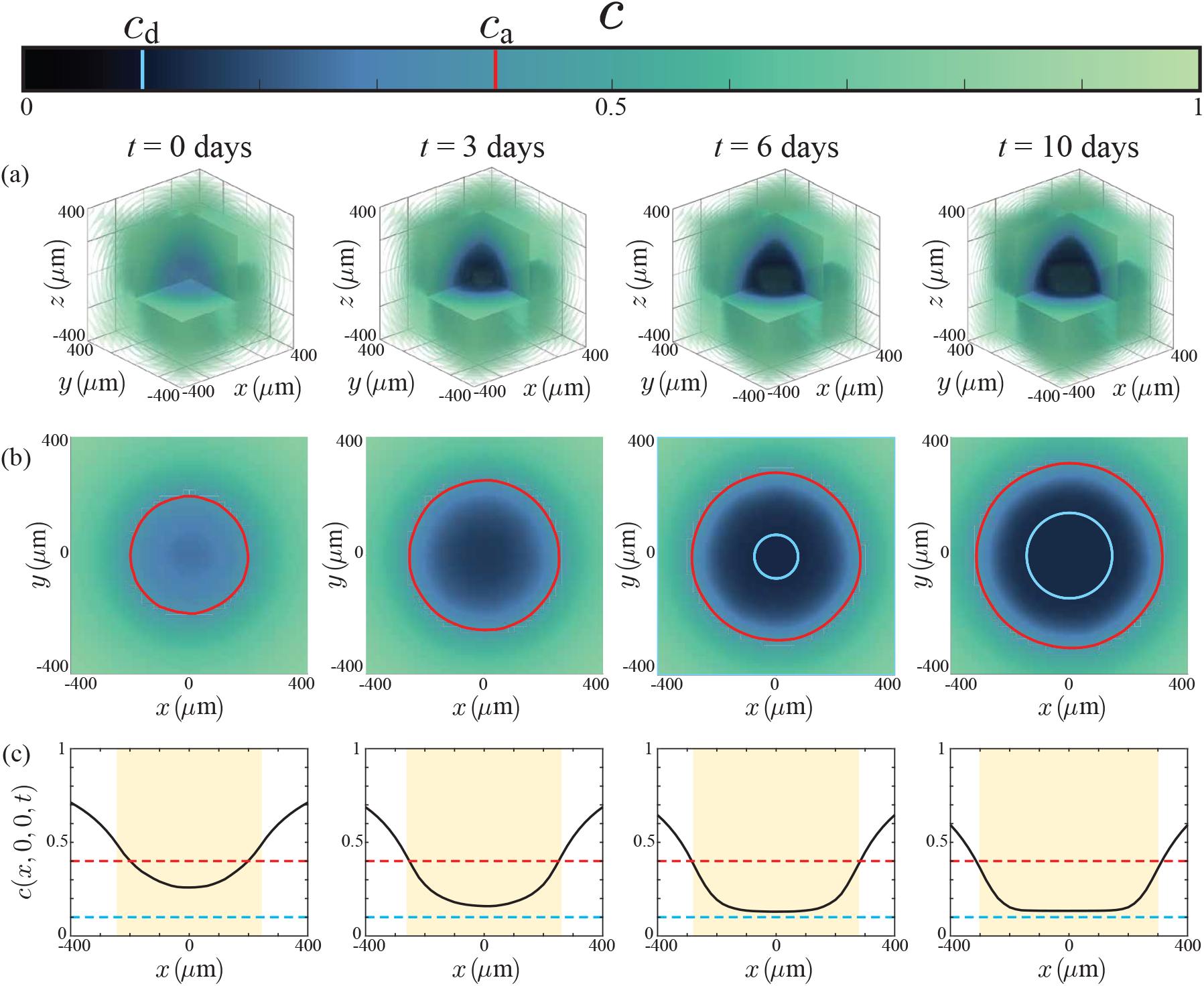
Nutrient concentration profiles (a) in three spatial dimensions, (b) at the equator *z* = 0, with the arrest critical level *c*_a_ shown in red, and the size of the necrotic region in white. (c) Nutrient profiles along the midline *y* = *z* = 0, where the shaded region represents the size of the spheroid, and the red and cyan lines are the critical levels for arrest and death, *c*_a_ and *c*_d_ respectively. The colourbar corresponds to the profiles in (a)–(b), and denotes the values *c*_a_ (red) and *c*_d_ (cyan).

While it is very difficult to measure the spatial and temporal distribution of diffusible nutrient experimentally in the growing spheroid, it is possible to indirectly examine our assumption that spatial and temporal differences in cell cycle status are partly driven by the availability of oxygen. Figure 6 shows a series of spheroids stained with pimonidazole and pimonidazole-detecting antibodies, which indicate hypoxia [40]. In this series of images, we see evidence of hypoxia staining in the central region of the spheroid at *t* = 0, with persistent hypoxia staining adjacent to the necrotic core at later times. These results support our hypothesis that spatial and temporal differences in nutrient availability correspond with spatial and temporal differences in cell cycle status, and in this case the pimonidazole staining suggests that oxygen availability plays a role in the development of heterogeneity within the growing population. While this observation is consistent with our IBM, it does not rule out the possibility of multiple diffusible signals acting in unison, and we will discuss this possibility later.

**Figure 6:**
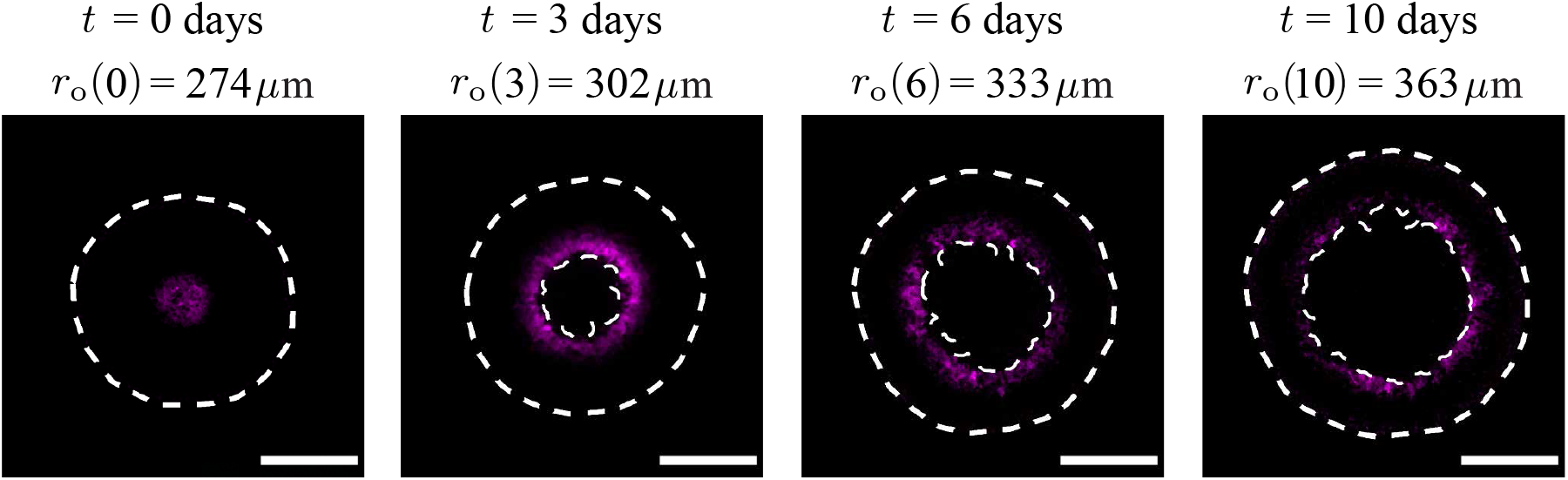
Spheroids stained for hypoxia at 0, 3, 6, and 10 days after spheroid formation, imaged at the spheroid equator. Hypoxia-positive staining fluoresces magenta, and white dashed lines denote *r*_o_(*t*) and *r*_n_(*t*), detected with image processing, to contextualise the regions of hypoxia. For clear visualisation, we label the outer radii of the spheroid with the corresponding days. Image intensity was adjusted for visual purposes, and scale bar corresponds to 200 *μ*m.

### 3.4. Role of variability

Experimental images (Figure 1, Figure 3, Figure 6) suggest that spheroid development is quite variable, as we see spheroids of slightly different diameters at the same time points. One of the limitations of relying on experimentation alone is that it can be difficult to quantify the importance of different sources of variability, whereas this can be assessed very simply with the IBM. For example, we can simulate multiple spheroids that start from precisely the same initial condition to quantify the variability that arises due to the stochastic growth process, or we can deliberately introduce variability into the initial composition of the spheroid to explore how this variability evolves during spheroid growth for a suite of simulated spheroids.

Simulation data in Figure 7a show the temporal evolution of various agent subpopulations, including the total number of living agents, dead agents, G1, eS, S/G2/M, and G1-arrested agents. Each profile shows the mean number of agents obtained by simulating 10 identically initialised spheroids with *r*_o_(0) = 245 *μ*m, which matches the average spheroid diameter at *t* = 0 days in the suite of *in vitro* experiments. The variability in these profiles is quantified by calculating the sample mean and sample standard deviation and shading the region corresponding to the sample mean plus or minus one sample standard deviation, and we see that, at this scale, the variability is barely noticeable. In contrast, results in Figure 7b show equivalent data from a suite of simulations where the initial density of agents in the spheroid is held constant, but the initial radius of the 10 simulated spheroids is deliberately varied to mimic the observed variability in our experiments. The initial radius in each simulation corresponds to one of 10 particular experimental measurements (Figure 7), with a sample mean of 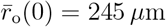 Comparing results in Figure 7a-b shows that the average population profiles are very similar, but the variability is strikingly different. This simple exercise shows that quantifying the variability in spheroid size at the beginning of the experiment is the key to understanding and predicting the variability in spheroid composition and size at the end of the experiment. We also see that 10 simulations is sufficient to observe the difference in variability between both test cases, where the spheroids start from identical initialisations or with induced variability. These simulation results are also consistent with our previous observations. For example, the *in vitro* spheroids in Figure 3 have *r*_o_(0) = 232 *μ*m and we see that it takes until *t* = 6 days for a clear necrotic core to form in the equatorial cross section. In contrast, the spheroid in Figure 6 is larger with *r*_o_(0) = 274 *μ*m and we see a clear necrotic core at *t* = 3 days. This highlights the importance of taking great care with measurements at the beginning of the experiment [36].

**Figure 7:**
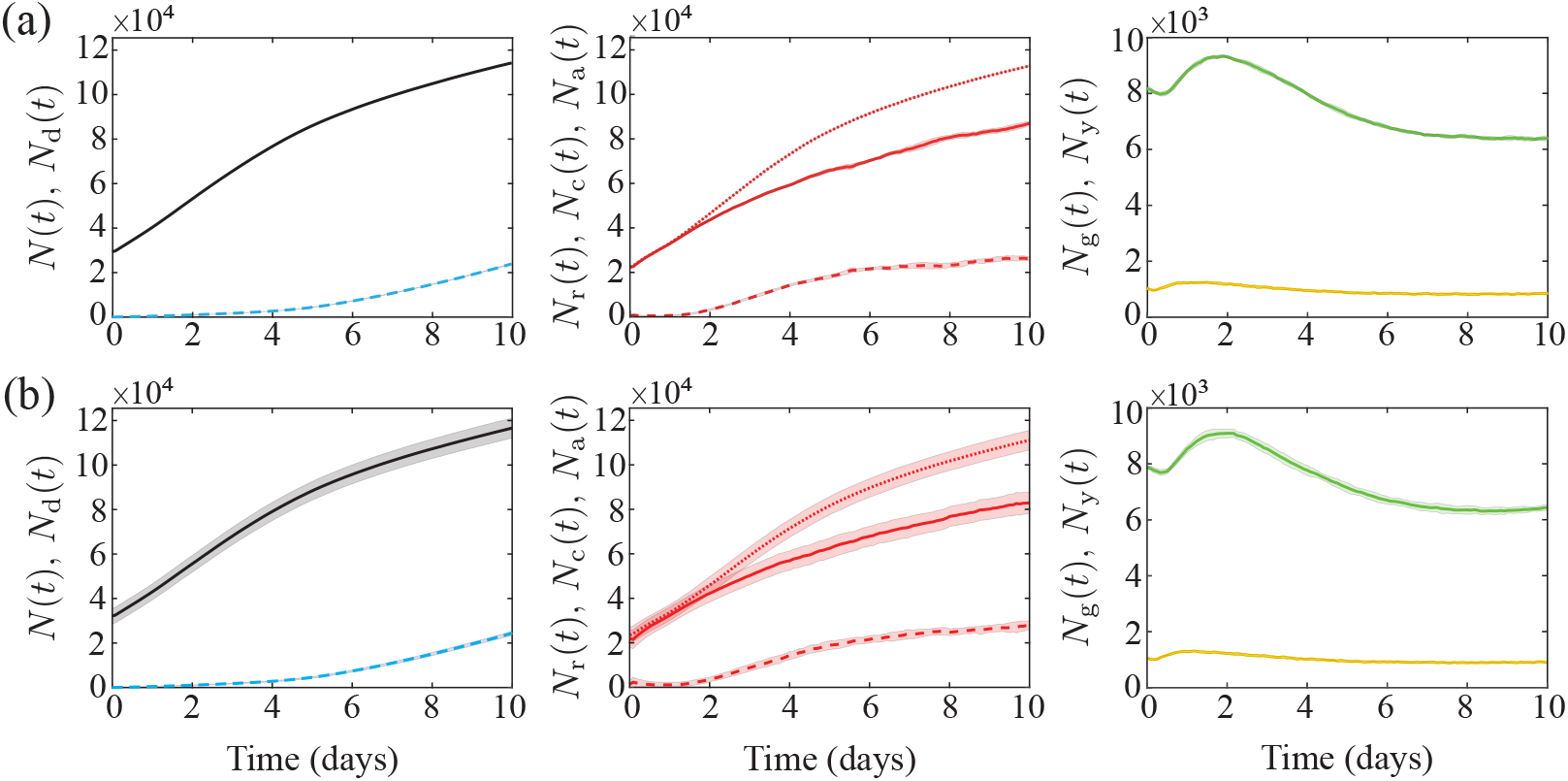
Modelling results for the population growth of different spheroid populations, averaged over 10 simulations with (a) identical initial conditions for each realisation and (b) introduced experimental variability in initial spheroid radius and population, with the agent density held constant and initial radius *r*_o_(*t*) ∈ [232.75, 235.47, 238.97, 242.19, 244.89, 247.76, 247.93, 251.23, 251.48, 260.13] *μ*m. In each row, left: living (black) and dead (cyan dashed) populations, *N*(*t*) and *N*_d_(*t*), respectively, centre: arrested red (dashed), cycling red (solid), and total red (dotted) populations, *N*_a_(*t*), *N*_c_(*t*), and *N*_r_(*t*), respectively, and right: yellow and green populations, *N*_y_(*t*) and *N*_g_(*t*), respectively. Shaded areas represent plus or minus one standard deviation. Initial subpopulations in each simulation in both (a) and (b) are variable, as initial cell cycle status is assigned randomly (Supplementary S7), and so the initial subpopulations in (b) also naturally vary with the overall initial population, *N*(0).

### 3.5. Quantitatively matching experimental and mathematical spheroids

Results in Figure 8 compare the temporal evolution of *r*_o_(*t*), *r*_a_(*t*), and *r*_n_(*t*), from our suite of experiments and simulations. The data in Figure 8 show the value in working with a stochastic model since the experimental measurements are quite variable, with estimates of *r*_a_(*t*) and *r*_n_(*t*) more variable than estimates of *r*_o_(*t*). This difference in variability is because we measure *r*_o_(*t*) automatically with an IncuCyte S3 every 6 hours. In contrast, measurements of *r*_a_(*t*) and *r*_n_(*t*) require manual harvesting, fixing, and imaging, and accordingly we report these measurements daily. Similarly to Section 3.4, we compare experimental results of average data in simulations with and without induced variability in the initial condition. The experiment-IBM comparison in Figure 8a corresponds to the case where we simulate 10 identically-prepared realisations of the IBM, where each simulated spheroid has the same initial radius *r*_o_(0) = 245 *μ*m, and we see that the average simulation results capture the average trends in the experimental measurements well, but the IBM simulations do not capture observed variability in the evolution of *r*_a_(*t*) or *r*_n_(*t*). In contrast, the experiment-IBM comparison in Figure 8b, where we deliberately mimic the experimental variability at *t* = 0, captures both the average experimental trends and variability in the experimental data quite well. Again, the difference between Figure 8a-b suggests that incorporating the initial variability in the experimental data is critical if we wish to capture the observed variability in the experiments.

**Figure 8:**
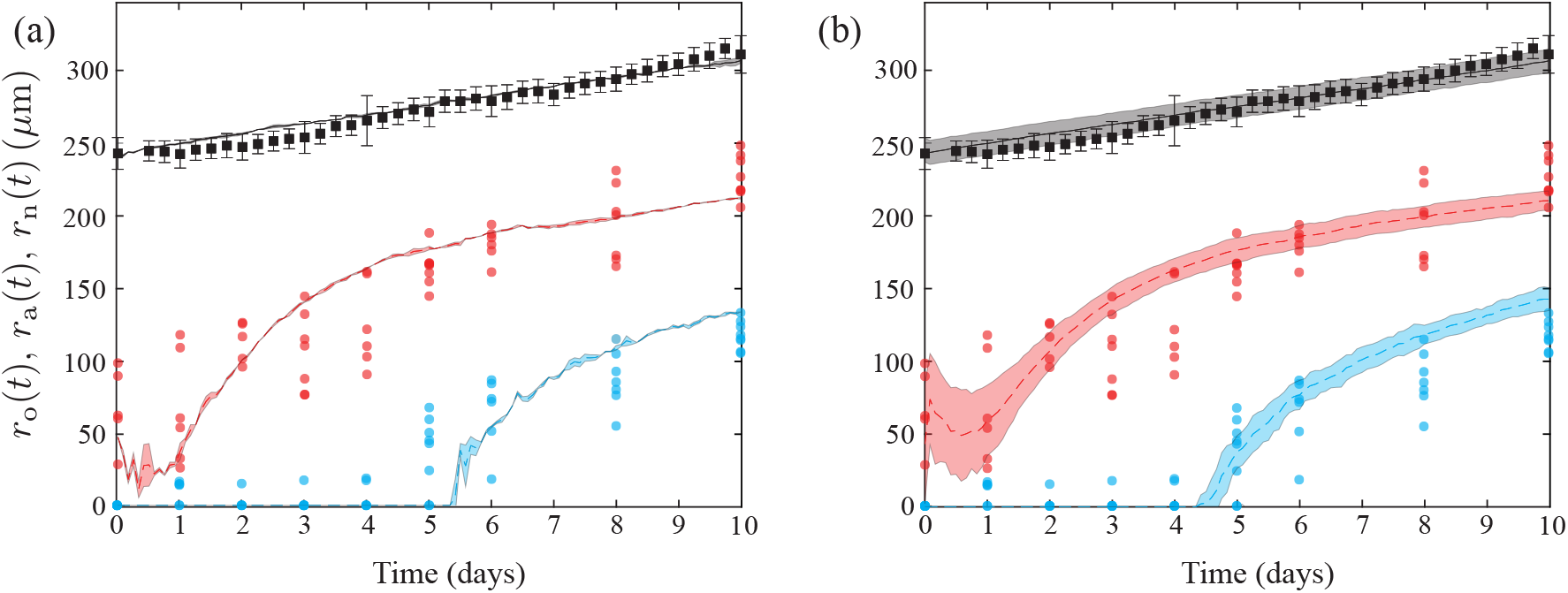
Comparison of computational estimates of *r*_o_(*t*) (black), *r*_a_(*t*) (red), and *r*_n_(*t*) (cyan) with experimental data. The experimental data (dots) are compared with (a) simulations with each run starting with an identical parameter set and (b) simulations with variations of the initial spheroid radius and population, with each initial radius selected from experimentally measured radii at *t* = 0 days and agent density kept constant. Computational results are the average of 10 simulations, and error regions represent plus or minus one standard deviation. The initial subpopulations vary in both (a) and (b), due to randomly assigning cell cycle status (Supplementary S7). In (b), we also naturally see higher variations in each subpopulation initially, due to explicitly including initial population variability, which in turn induces variability in *r*_a_(0).

Interestingly, our experimental data in Figure 8 suggest that we have an approximately linear increase in *r*_o_(*t*) over time, whereas the development of the internal structure is more complicated. The initial arrested radius decreases for the first day before growing rapidly, and we do not see the formation of a necrotic core until approximately *t* = 4 days. While our IBM-experimental comparison in Figure 8 suggests that the IBM can quantitatively capture experimental trends, we have obtained this match with a careful choice of parameters without undertaking a more rigorous parameter estimation exercise [41].

## 4. Conclusions and Future Work

In this work we develop a novel IBM that can simulate 4D tumour spheroid experiments with explicit cell cycle labels. IBM simulations reveal that we can successfully reproduce qualitative and quantitative patterns of spatial and temporal differences in cell cycle status that we observe in *in vitro* experiments. This heterogeneity is driven by spatial and temporal variations in nutrient availability, which we model using a reaction-diffusion equation coupled to the IBM.

An important advantage of the IBM is our ability to extract and describe measurements that are difficult to obtain *in vitro*. In particular, we show how to visualise both the growing populations within the spheroid together with the spatial patterns of nutrient concentration over time within the growing spheroid. Furthermore, the IBM makes it very simple to explore how various features contribute to the overall variability in spheroid development, and we find that relatively small variations in the initial size of the spheroid lead to relatively pronounced differences in spheroid size and composition at later times [36]. We conclude our investigation by showing that we can quantitatively match the spatial and temporal development of a series of *in vitro* 4D spheroids using the WM793B human primary melanoma cell line with a careful choice of parameters. We anticipate that tumour spheroids formed with different cell lines will be able to be simulated with our IBM, but will require different parameter values.

Overall, our modelling philosophy is always to work with the simplest possible mechanisms required to capture our experimental observations. Naturally, this means that there are many ways that the IBM can be extended. For example, here we make the simple assumption that spheroid growth is regulated by a single diffusible nutrient, which seems appropriate for our data. If, however, experiments show that it is important to consider multiple nutrients in unison, our IBM framework can be extended to deal with this. Similarly, we have focused on spheroid growth commencing with a spherically symmetric initial condition which is consistent with our experiments. This assumption can be relaxed in the present model simply by specifying a different arrangement of agents at *t* = 0. Another point that could be revisited is that we implement the simplest possible cell migration mechanism where the direction of motion is random. While this assumption appears reasonable for our data, it is possible to bias the migration in response to either the nutrient concentration, the gradient of the nutrient concentration, or the density of agents. Each of these potential extensions could be incorporated into our IBM framework and increase the biological fidelity of the model. However, here we caution against this approach since these mechanisms also increase the number of parameters required for simulation. To minimise issues with parameter identifiability, we prefer to work with a minimal model [41]. If, however, future experimental measurements indicate that our minimal assumptions need to be revised, our IBM framework is sufficiently flexible to incorporate such extensions, if warranted. Another option for future refinement is to conduct a more thorough parameter estimation exercise. Here we carefully chose parameters that appear to match our data, but future analysis could include a more rigorous assessment of parameter estimation, and we leave this for future consideration.

## Supporting information

Additional Results and Discussion

## Acknowledgements

MJS, NKH, and MJP are supported by the Australian Research Council (DP200100177). We appreciate computational resources of the QUT High Performance Computing support group.

## References

[1] Nunes AS, Barris AS, Costa EC, Moreira AF, Correia IJ, 2018. 3D tumor spheroids as in vitro models to mimic in vivo human solid tumors resistance to therapeutic drugs. Biotechnology and Bioengineering, 116:206–226. doi: 10.1002/bit.26845.

[2] Lazzari G, Couvreur P, Mura S, 2017. Multicellular tumor spheroids: a relevant 3D model for the in vitro preclinical investigation of polymer nanomedicines. Polymer Chemistry, 8:4947–4969. doi: 10.1039/C7PY00559H.

[3] Spoerri L, Beaumont KA, Anfosso A, Haass NK, 2017. Real-time cell cycle imaging in a 3D cell culture model of melanoma. Methods in Molecular Biology, 1612:401–416. doi: 10.1007/978-1-4939-7021-629.

[4] Haass NK, Beaumont KA, Hill DS, Anfosso A, Mrass P, Munoz MA, Kinjyo I, Weninger W, 2014. Real-time cell cycle imaging during melanoma growth, invasion, and drug response. Pigment Cell & Melanoma Research, 27:764–776. doi: 10.1111/pcmr.12274.

[5] Mehta G, Hsiao AY, Ingram M, Luker GD, Takayama S, 2012. Opportunities and challenges for use of tumor spheroids as models to test drug delivery and efficacy. Journal of Controlled Release, 164:192–204. doi: 10.1016/j.jconrel.2012.04.045.

[6] Beaumont KA, Mohana-Kumaran N, Haass NK, 2014. Modelling melanoma in vitro and in vivo. Healthcare, 2:27–46. doi: 10.3390/healthcare2010027.

[7] Sakaue-Sawano A, Kurokawa H, Morimura T, Hanyu A, Hama H, Osawa H, Kashiwagi S, Fukami K, Miyata T, Miyoshi H, et al., 2008. Visualizing spatiotemporal dynamics of multicellular cell-cycle progression. Cell, 132:487–498. doi: 10.1016/j.cell.2007.12.033.

[8] Vittadello ST, McCue SW, Gunasingh G, Haass NK, Simpson MJ, 2019. Mathematical models incorporating a multi-stage cell cycle replicate normally-hidden inherent synchronoization in cell proliferation. Journal of the Royal Society Interface, 16:20190382. doi: 10.1098/rsif.2019.0382.

[9] Jin W, Spoerri L, Haass NK, Simpson MJ, 2021. Mathematical model of tumour spheroid experiments with real-time cell cycle imaging. Bulletin of Mathematical Biology, 83:1–23. doi: 10.1007/s11538-021-00878-4.

[10] Greenspan HP, 1972. Models for the growth of a solid tumor by diffusion. Studies in Applied Mathematics, 51:317–340. doi: 10.1002/sapm1972514317.

[11] McElwain DLS, Ponzo PJ, 1977. A model for the growth of a solid tumor with non-uniform oxygen consumption. Mathematical Biosciences, 35:267–279. doi: 10.1016/0025-5564(77)90028-1.

[12] Ward JP, King JR, 1997. Mathematical modelling of avascular-tumour growth. Mathematical Medicine and Biology, 14:39–69. doi: 10.1093/imammb/14.1.39.

[13] Landman KA, Please CP, 2001. Tumour dynamics and necrosis: surface tension and stability. Mathematical Medicine and Biology, 18:131–158. doi: 10.1093/imammb/18.2.131.

[14] Byrne HM, Chaplain MAJ, 1995. Growth of nonnecrotic tumors in the presence and absence of inhibitors. Mathematical Biosciences, 130:151–181. doi: 10.1016/0025-5564(94)00117-3.

[15] Byrne HM, King JR, McElwain DLS, Preziosi L, 2003. A two-phase model of solid tumour growth. Applied Mathematics Letters, 16:567–573. doi: 10.1016/S0893-9659(03)00038-7.

[16] Leedale J, Herrmann A, Bagnall J, Fercher A, Papkovsky D, Sée V, Bearon RN, 2014. Modeling the dynamics of hypoxia inducible factor-1α(HIF-1α) within single cells and 3D cell culture systems. Mathematical Biosciences, 258:33–43. doi: 10.1016/j.mbs.2014.09.007.

[17] Grimes DR, Kelly C, Bloch K, Partridge M, 2014. A method for estimating the oxygen consumption rate in multicellular tumour spheroids. Journal of the Royal Society Interface, 11:20131124. doi: 10.1098/rsif.2013.1124.

[18] Browning AP, Sharp JA, Murphy RJ, Gunasingh G, Lawson B, Burrage K, Haass NK, Simpson MJ, 2021. Quantitative analysis of tumour spheroid structure. bioRxiv. doi: 10.1101/2021.08.05.455334. To appear, eLife.

[19] Codling EA, Plank MJ, Benhamou S, 2008. Random walk models in biology. Journal of the Royal Society Interface, 5:813–834. doi: 10.1098/rsif.2008.0014.

[20] Mallet DG, De Pillis LG, 2006. A cellular automata model of tumor-immune system interactions. Journal of Theoretical Biology, 239:334–350. doi: 10.1016/j.jtbi.2005.08.002.

[21] Browning AP, McCue SW, Binny RN, Plank MJ, Shah ET, Simpson MJ, 2018. Inferring parameters for a lattice-free model of cell migration and proliferation using experimental data. Journal of Theoretical Biology, 437:251–260. doi: 10.1016/j.jtbi.2017.10.032.

[22] Browning AP, Jin W, Plank MJ, Simpson MJ, 2020. Identifying density-dependent interactions in collective cell behaviour. Journal of the Royal Society Interface, 17:20200143. doi: 10.1098/rsif.2020.0143.

[23] Carr MJ, Simpson MJ, Drovandi C, 2021. Estimating parameters of a stochastic cell invasion model with fluorescent cell cycle labelling using approximate Bayesian computation. Journal of the Royal Society Interface, 18:20210362. doi: 10.1098/rsif.2021.0362.

[24] Mao X, McManaway S, Jaiswal JK, Patel PB, Wilson WR, Hicks KO, Bogle G, 2018. An agent-based model for drug-radiation interactions in the tumour microenvironment: hypoxia-activated prodrug SN30000 in multicellular tumour spheroids. PLoS Computational Biology, 14:e1006469. doi: 10.1371/journal.pcbi.1006469.

[25] Bull JA, Mech F, Quaiser T, Waters SL, Byrne HM, 2020. Mathematical modelling reveals cellular dynamics within tumour spheroids. PLoS Computatinal Biology, 16:e1007961. doi: 10.1371/journal.pcbi.1007961.

[26] Hoek KS, Schlegel NC, Brafford P, Sucker A, Ugurel S, Kumar R, Weber BL, Nathanson KL, Phillips DJ, Herlyn M, et al., 2006. Metastatic potential of melanomas defined by specific gene expression profiles with no BRAF signature. Pigment Cell Research, 19:290–302. doi: 10.1111/j.1600-0749.2006.00322.x.

[27] Smalley KSM, Contractor R, Haass NK, Kulp AN, Atilla-Gokcumen GE, Williams DS, Bregman H, Flaherty KT, Soengas MS, Meggers E, et al., 2007. An organometallic protein kinase inhibitor pharmacologically activates p53 and induces apoptosis in human melanoma cells. Cancer Research, 67:209–217. doi: 10.1158/0008-5472.CAN-06-1538.

[28] Smalley KSM, Contractor R, Haass NK, Nathanson KL, Medina CA, T. Fk, Herlyn M, 2007. Ki67 expression levels are a better marker of reduced melanoma growth following MEK inhibitor treatment than phospho-ERK levels. British Journal of Cancer, 96:445–449. doi: 10.1038/sj.bjc.6603596.

[29] Spoerri L, Gunasingh G, Haass NK, 2021. Fluorescence-based quantitative and spatial analysis of tumour spheroids: a proposed tool to predict patient-specific therapy response. Frontiers in Digital Health, 3:1–19. doi: 10.3389/fdgth.2021.668390.

[30] Cold Spring Harbor Laboratory Press, 2018. Antibody dilution buffer (Abdil) protocol. doi: 10.1101/pdb.rec103978 (Accessed: November 2021).

[31] Browning AP, Murphy RJ, 2021. Image processing algorithm to identify structure of tumour spheroids with cell cycle labelling. Zenodo. doi: 10.5281/zenodo.5121093.

[32] Gillespie DT, 1977. Exact stochastic simulation of coupled chemical reactions. The Journal of Physical Chemistry, 81:2340–2361. doi: 10.1021/j100540a008.

[33] Weisstein EW, 2021. Sphere point picking. mathworld – A Wolfram web resource. https://mathworld.wolfram.com/SpherePointPicking.html. (Accessed: November 2021).

[34] Simpson MJ, Landman KA, Hughes BD, 2010. Cell invasion with proliferation mechanisms motivated by time-lapse data. Physica A: Statistical Mechanics and its Applications, 389:3779–3790. doi: 10.1016/j.physa.2010.05.020.

[35] Treloar KK, Simpson MJ, 2013. Sensitivity of edge detection methods for quantifying cell migration assays. PLoS One, 8:e67389–e67389. doi: 10.1371/journal.pone.0067389.

[36] Murphy RJ, Browning AP, Gunasingh G, Haass NK, Simpson MJ, 2021. Designing and interpreting 4D tumour spheroid experiments. bioRxiv. doi: 10.1101/2021.08.18.456910.

[37] Spoerri L, Tonnessen-Murray CA, Gunasingh G, Hill DS, Beaumont KA, Jurek RJ, Chauhan J, Vanwalleghem GC, Fane ME, Daignault-Mill SM, et al., 2021. Phenotypic melanoma heterogeneity is regulated through cell-matrix interaction-dependent changes in tumor microarchitecture. bioRxiv preprint. doi: 10.1101/2020.06.09.141747.

[38] Miniaev MV, Belyakova MB, Kostiuk NV, Leshchenko DV, Fedotova TA, 2013. Non-obvious problems in Clark electrode application at elevated temperature and ways of their elimination. Journal of Analytical Methods in Chemistry, 2013:249752. doi: 10.1155/2013/249752.

[39] Langan LM, Dodd NJF, Owen SF, Purcell WM, Jackson SK, Jha AN, 2016. Direct measurements of oxygen gradients in spheroid culture system using electrion parametric resonance oximetry. PLoS One, 11:e0149492. doi: 10.1371/journal.pone.0149492.

[40] Varia MA, Calkins-Adams DP, Rinker LH, Kennedy AS, Novotny DB, Fowler WC, J., Raleigh JA, 1998. Pimonidazole: a novel hypoxia marker for complementary study of tumor hypoxia and cell proliferation in cervical carcinoma. Gynecologic Oncology, 71:270–277. doi: 10.1006/gyno.1998.5163.

[41] Simpson MJ, Baker RE, Vittadello ST, Maclaren OJ, 2020. Practical parameter identifiability for spatio-temporal models of cell invasion. Journal of the Royal Society Interface, 17:20200055. doi: 10.1098/rsif.2020.0055.

