## Additional Results and Discussion for "A stochastic mathematical model of 4D tumour spheroids with real-time fluorescent cell cycle labelling"

**S4 Estimation of cell diameter** **16**

**S5 Estimation of cell cycle progression rates** **17**

**S6 Initial cell number** **18**

**S7 Initialising the IBM** **20**

**S8 Simulation algorithms** **22**

**S9 Calculation of agent density profiles** **24**

†These authors contributed equally to this work.

### S1. Image processing

The image processing algorithm for experimental images uses the algorithm presented and described in [1]. Here, we describe the procedure used to prepare the synthetic data from the IBM for estimates of $r_{\mathrm{o}}(t)$ and $r_{\mathrm{n}}(t)$, and visualise the results in Figure S1. We perform the image analysis on synthetic data in MATLAB as follows:

1. An image of $(L+1)\times(L+1)$ pixels is created, so that each pixel is one micrometre in length,and all agents within 18 $\mu$m of $z = 0$ have their $x$ and $y$ coordinates recorded,
2. Living agents from the synthetic dataset are rounded to their nearest integer location with `round` and placed at the corresponding pixel in the image with `sub2ind`,
3. Each agent is increased in size to a diameter of $\sigma$ pixels,
4. Components of the binarised image with a connected area less than a threshold value $A$ are removed by area opening with `bwareaopen`, where $A$ is sufficiently large so that only the contiguous spheroid remains,
5. Remaining holes are filled (`imfill`) and remaining peripheral segments disconnected from the main body are removed (`imclearborder`),
6. The area enclosed by the boundary is calculated with `regionprops`, and $r_{\mathrm{o}}(t)$ is estimated by assuming the region is circular, and calculating the radius of a circle with an equivalent area,
7. A mask of dead agents is created according to steps 1–3,
8. The intersection between the dead agent mask and the space left unfilled by the living agents before performing step 5 is found,
9. If the radius of a circle with equivalent area to this intersection is smaller than $2\Delta_{\mathrm{c}}$ (too small to be meaningful in the context of a necrotic core), set the necrotic radius, $r_{\mathrm{n}}(t) = 0$,
10. Otherwise, $r_{\mathrm{n}}(t)$ is set as the radius of a circle with equivalent area to the filled space of the dead agent mask.

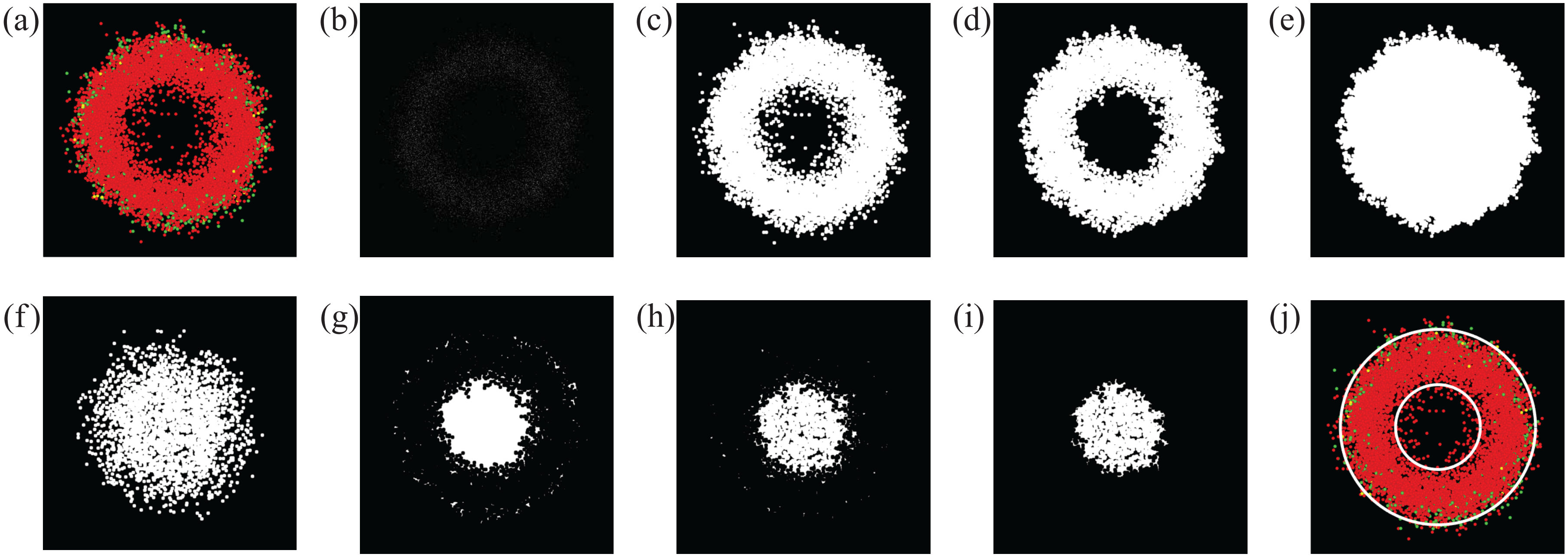

**Figure S1:** Example of applying the radius estimate image processing algorithm to a synthetic dataset. (a) A 2D slice at the equator of the spheroid, (b) the agents are mapped into a single pixel each at their equivalent positions on a binary, $(L+1)\times(L+1)$ image. The agent size is then increased, as shown in (c), to better represent their equivalent physical representation, before area opening (d) and filling the remaining space (e). The radius of a circle with equivalent area to the filled space is calculated, and used as the estimate for the outer radius, $r_{\mathrm{o}}(t)$. (f) The mask of dead agents is created, along with (g) a mask of the empty space inside the spheroid in (d). (h) The intersection between the dead agent mask and the empty space, and (i) the clearing of (h) to only the largest continuous area. If this region is large enough, the necrotic region is taken to be the largest area in (g), and its radius, $r_{\mathrm{n}}(t)$, is the radius of a circle with equivalent area. (j) The estimates for $r_{\mathrm{o}}(t)$ and $r_{\mathrm{n}}(t)$ are shown plotted over the original spheroid slice.

Similarly to the estimates of $r_{\mathrm{o}}(t)$ and $r_{\mathrm{n}}(t)$, we estimate $r_{\mathrm{a}}(t)$ with MATLAB, and the procedure is designed to mimic the procedure used for estimating the arrested radius in experimental images, outlined in [1]. Visualisation of this process is given in Figure S2. We prepare the synthetic data as follows:

1. Each green agent from the synthetic dataset at time $t$ is converted from its 3D Cartesian coordinate, $\mathbf{x}_n$, to its radial coordinate, $r_n$, with the transformation

$$r_n = ||\mathbf{x}_n - \bar{\mathbf{x}}(t)||, \qquad n = 1, \ldots, N_g(t), \tag{S1}$$

   where $N_g(t)$ is the number of green agents at time $t$ and $\bar{\mathbf{x}}(t)$ is the mean location of all agents in the spheroid at time $t$ (Figure S2a), and $||\cdot||$ is the Euclidean length,

2. The green agents are organised into a distribution of radial position, $G(r_l)$, where $r_l$ are the centres of the bins with binwidth $w = 20$ $\mu$m, so that $l = 0, \ldots, \texttt{round}\,(r_{\mathrm{o}}(t)/w)$. We set $l_{\max} = \texttt{round}\,(r_{\mathrm{o}}(t)/w)$ as this maximum index. The binwidth $w = 20\,\mu$m is chosen to be sufficiently small to best identify spatial information about the radial distribution of green agents, but also large enough to avoid fluctuations in the distribution [2],

3. The agent density, $g(r_l)$, is calculated by dividing the agent count distribution, $G(r_l)$, by the radial volume,

$$g(r_l) = \frac{3G(r_l)}{4\pi\left(\left(r_l + \dfrac{w}{2}\right)^3 - \left(r_l - \dfrac{w}{2}\right)^3\right)}, \qquad l = 0, \ldots, l_{\max}, \tag{S2}$$

4. The agent density is then normalised, such that

$$\hat{g}(r_l) = \frac{g(r_l)}{\texttt{max}(g(r_l))}, \qquad l = 0, \ldots, l_{\max}, \tag{S3}$$

   where the $\texttt{max}(g(r_l))$ is the maximum of the green agent density at time $t$,

5. A Gompertz function,

$$\hat{g}_{\mathrm{G}}(r) = \gamma_1 \exp\left(-\exp\left(\gamma_2\left(\gamma_3 - r\right)\right)\right), \tag{S4}$$

is fit to the normalised agent density distribution, $\hat{g}(r_l)$, on the domain $r \in [0, \arg\max(\hat{g}(r_l))]$ using the method of least squares to create a smoothed normalised green agent density as a function of radial position, $\hat{g}_{\mathrm{G}}(r)$ (Figure S2b). The values $\gamma_1$, $\gamma_2$, $\gamma_3$ are the parameters fit by the method of least squares. This choice of domain, from $r = 0$ to $r = \arg\max(\hat{g}(r_l))$, is used because it generates a more accurate fit of $\hat{g}_G(r)$ to $\hat{g}(r_l)$ in the region of interest, due to the shapes of the Gompertz function and normalised agent density function, $\hat{g}(r_l)$,

6. Adapting the method in [1], if $\hat{g}_{\mathrm{G}}(0) > g_{\mathrm{a}}$, then $r_{\mathrm{a}}(t) = 0$. Otherwise, $r_{\mathrm{a}}(t)$ is defined such that $\hat{g}_{\mathrm{G}}(r_{\mathrm{a}}(t)) = g_{\mathrm{a}}$. We use the tunable threshold parameter $g_{\mathrm{a}} = 0.2$, so that the arrested region is considered to be the boundary where the smoothed relative green agent density function surpasses 0.2.

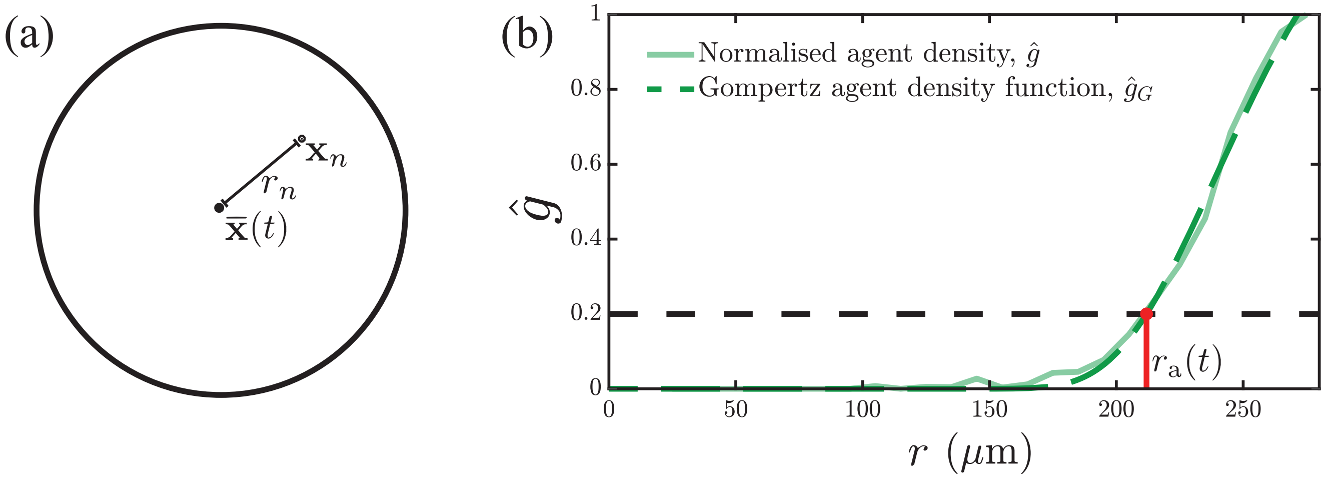

**Figure S2:** Example of how the arrested radius is estimated with synthetic data. (a) The 3D Cartesian position of a green agent, $\mathbf{x}_n$, is converted into a radial coordinate, $r_n$, by calculating its Euclidean distance from the mean position of all agents in the spheroid, $\bar{\mathbf{x}}(t)$. (b) An example of the normalised green agent density distribution, $\hat{g}(r_l)$, and the Gompertz function of smoothed agent density, $\hat{g}_G(r)$, for a spheroid at late time. The arrested radius, $r_{\mathrm{a}}(t)$ is calculated as the value of $r$ for which $\hat{g}_G(r) = g_{\mathrm{a}}$, where $g_{\mathrm{a}} = 0.2$.

### S2. Benchmarking the 3D numerical partial differential equation solution

To ensure our spatial discretisation of the 3D partial differential equation (PDE) model of the nutrient profile is accurate, we first solved a number of simpler test cases with exact solutions. To this end, we consider an infinite domain 3D diffusion-decay problem where some mass of diffusing substance, $S$, is placed at the origin at $t = 0$ [3]. The governing equation is

$$\frac{\partial c}{\partial t} = D\nabla^2 c - \kappa c, \tag{S5}$$

with diffusivity $D > 0$ and decay rate $\kappa$. The exact solution is,

$$c(r,t) = \frac{S}{\left(2\sqrt{\pi D t}\right)^3} \exp\left(-\frac{r^2}{4Dt} - \kappa t\right), \tag{S6}$$

where $r = \sqrt{x^2 + y^2 + z^2}$ [3].

To test the accuracy of our numerical method, we consider this problem on a truncated finite domain. In this case we consider a cube of side length $L = 20$, with the origin at the centre of the cube. We discretise the truncated domain using a uniform mesh with node spacing $h = 0.2$ so that our mesh contains $101^3$ nodes. In our numerical simulations, we impose homogeneous Dirichlet boundary conditions along all boundaries. The numerical solution is obtained by discretising the flux and source terms in Equation (S5) on the finite volume mesh in exactly the same way as for the nutrient model in the main document. This leads to a system of coupled ordinary differential equations that we solve in time using MATLAB's `ode45` solver [4]. Results in Figure S3 compare exact and numerical solutions along the $y = z = 0$ midline for $x \in [-5, 5]$. In Figure S3a we consider a conservative problem where $\kappa = 0$, and in Figure S3b we consider a diffusion-decay problem with $\kappa > 0$. In all cases the numerical solution matches the exact solution well, giving us confidence in our spatial discretisation method.

We set the total amount of mass diffusing through the domain at $S = 100$ and use diffusivity $D = 1$, and consider two decay parameters, $\kappa = 0$ (diffusion without a source term) and $\kappa = 0.3$. In Figure S3, we compare exact and numerical solutions at $t = 1$, $t = 1.5$, and $t = 2$. Due to the radial symmetry of our solution, we plot the numerical and exact solutions along the $y = z = 0$ midline for $x \in [-5, 5]$, which gives better clarity in our comparison.

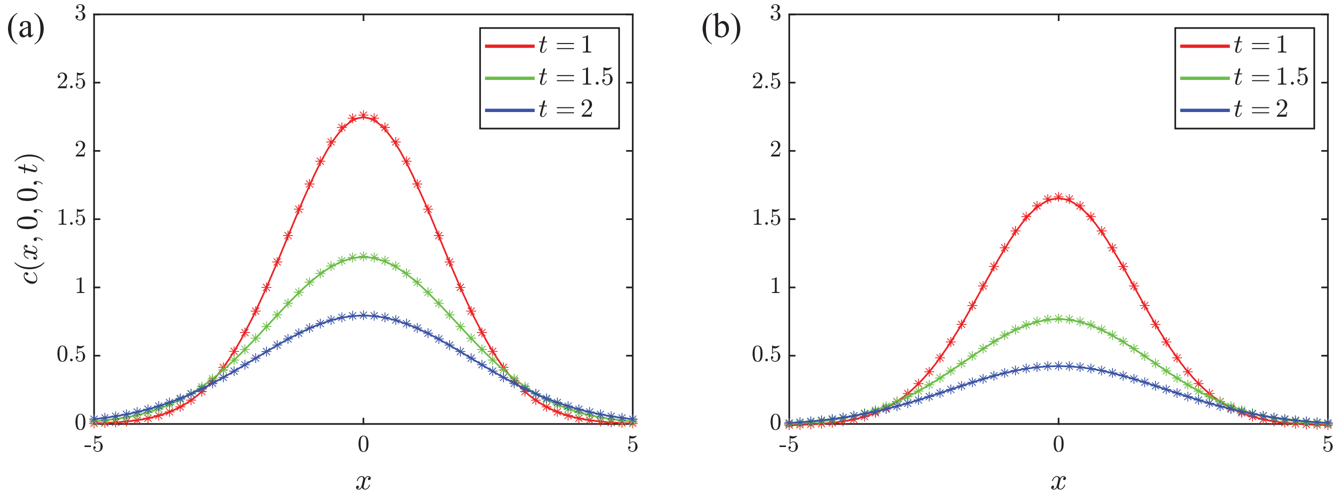

**Figure S3:** Comparison of exact (solid) and numerical (stars) solutions of Equation (S5). All results correspond to $S = 100$ and $D = 1$, with $h = 0.2$. Results in (a) are for $\kappa = 0$, whereas results in (b) are for $\kappa = 0.3$.

### S3. Numerical method and parameters

Here we provide the discretisation for the numerical method and additional evidence to justify our choices of the numerical parameters chosen for the simulation. In summary, these choices include tolerances for the numerical solution of the linear system associated with solving Equation (**??**); the size of the domain; the time duration between solving Equation (**??**); and the spatial discretisation required to solve Equation (**??**).

#### *S3.1. Numerical discretisation*

We solve Equation (**??**) by discretising the domain, $\Omega$, into a uniform structured finite volume mesh with cubic finite volumes, consisting of $I^3$ equally-spaced nodes with node spacing $h\,[\mu\text{m}]$. We approximate the cell density $v(\mathbf{x},t)$ in Equation (**??**) by the agent density, $N_{i,j,k}/h^3$, around node $(x_i, y_j, z_k)$, where $N_{i,j,k}$ [cells] is the the number of agents inside the node's control volume, and $h = L/(I-1)$. The discretised 3D finite volume equation is

$$h\left(c_{i-1,j,k} + c_{i,j-1,k} + c_{i,j,k-1} + c_{i+1,j,k} + c_{i,j+1,k} + c_{i,j,k+1} - 6c_{i,j,k}\right) - \alpha c_{i,j,k} N_{i,j,k} = 0, \qquad \text{(S7)}$$

for the internal nodes, and $c_{i,j,k} = 1$ at boundary nodes. Assembling these discrete equations gives a linear system that we solve numerically,

$$\mathbf{A}\mathbf{c} = \mathbf{b}, \qquad \text{(S8)}$$

where $\mathbf{A} \in \mathbb{R}^{I^3 \times I^3}$ is the coefficient matrix and $\mathbf{b} \in \mathbb{R}^{I^3}$ is the right-hand side vector for Equation (S7), and $\mathbf{c} \in \mathbb{R}^{I^3}$ is the solution.

#### *S3.2. GMRES tolerance*

To solve Equation (S8), we use the generalised minimal residual method (GMRES) in MATLAB [5]. Initially, we solve the linear system by assuming that $c = 1$ at all nodes and specify a strict relative tolerance $\varepsilon$ given by,

$$\varepsilon = \frac{||\mathbf{b} - \mathbf{A}\mathbf{c}||}{||\mathbf{b}||}, \qquad \text{(S9)}$$

where $||\cdot||$ is the 2-norm. Once this initial solution is obtained using a strict tolerance, we find that all subsequent solutions of Equation (S7) can be obtained with a larger tolerance since we have an improved estimate of the solution to initialise the iterative solver.

To determine the first tolerance required to solve the linear system, $\varepsilon_1$, we suppose that $c = 1$ at all nodes on a mesh with $I^3 = 201^3$ equally–spaced nodes (Section S3.4) and solve the resulting system with a range of $\varepsilon_1$ using MATLAB's `gmres` function. Table S1 compares the computation time as a function of $\varepsilon_1$, and Figure S4 compares the resulting nutrient profile solutions along the midline of the domain, $y = z = 0$.

**Table S1:** Runtimes for the initial solution to Equation (S8). Solution runtimes were achieved with high performance computing, using four 64 bit Intel Xeon core processors per simulation.

| $\varepsilon_1$ | Runtime (min.) |
|---|---|
| $1 \times 10^{-6}$ | 0.8 |
| $1 \times 10^{-7}$ | 13.0 |
| $1 \times 10^{-8}$ | 19.3 |
| $1 \times 10^{-9}$ | 24.4 |

Approaches utilising other, more specified initial guesses were also performed, but showed no improvement on runtimes for the initial solution. Since we see that the solutions for $\varepsilon_1 = 1 \times 10^{-8}$ and $1 \times 10^{-9}$ are visually indistinguishable at this scale, we always work with $\varepsilon_1 = 1 \times 10^{-8}$.

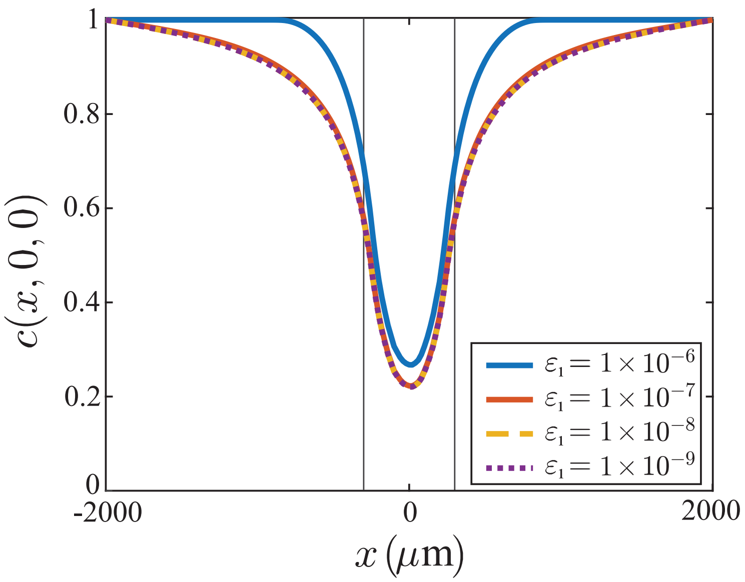

**Figure S4:** Comparison of steady-state solutions to Equation (S7) with different values of $\varepsilon_1$. Solutions with $\varepsilon_1 \leq 1 \times 10^{-8}$ are entirely visually indistinguishable, and those for a tolerance of $\varepsilon_1 = 1 \times 10^{-7}$ or lower visually match in the region occupied by the spheroid at $t = 0$ days, denoted by the vertical lines.

To update the nutrient distribution subsequently, we find that we do not need such a strict tolerance choice for subsequent solutions of Equation (S7), $\varepsilon_2$, as we have an improved estimate for the nutrient profile from the initial solution. To improve the estimate for the initial guess, we solve Equation (S7) on a relatively coarse mesh of $I^3 = 51^3$ nodes and solve the linear system using MATLAB's left matrix division (backslash) operator [6], and then interpolate this coarse solution onto the finer $I^3 = 201^3$ mesh, which serves as an initial guess for the next GMRES solution that can be obtained with a more relaxed tolerance. Table S2 compares the computation time between subsequent solutions as a function of $\varepsilon_2$.

**Table S2:** Runtimes for subsequent solutions to Equation (S8). Solution runtimes were achieved with high performance computing, using four 64 bit Intel Xeon core processors per simulation.

| $\varepsilon_2$ | Runtime (min.) |
|---|---|
| $1 \times 10^{-6}$ | 0.15 |
| $1 \times 10^{-7}$ | 0.95 |
| $1 \times 10^{-8}$ | 10.1 |
| $1 \times 10^{-9}$ | 20.6 |

Results in Figure S5 show $c(x, y, z)$ plotted along the midline $y = z = 0$ for one of these subsequent solutions for various choices of tolerances. Since we see that the solutions are all largely indistinguishable, and more restrictive tolerances are associated with significant increases in runtime with insignificant improvement in accuracy, we set $\varepsilon_2 = 1 \times 10^{-6}$ for all remaining calls to `gmres`.

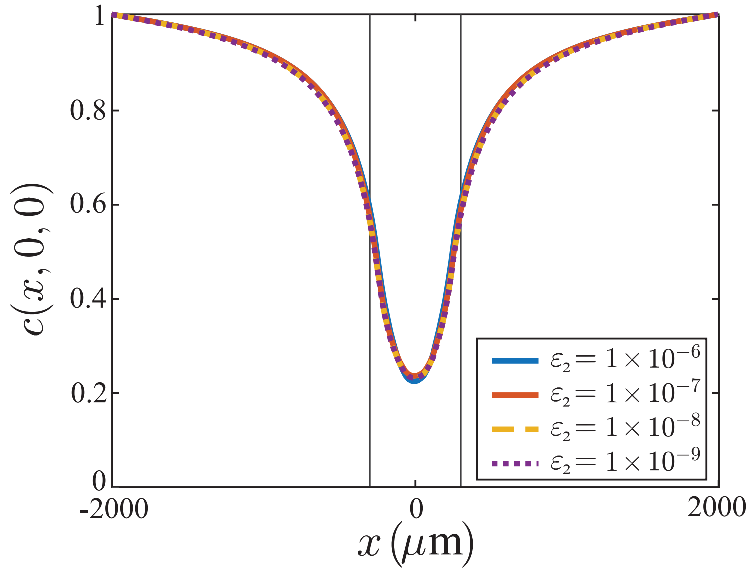

**Figure S5:** Comparison of GMRES solutions to Equation (S7) with different values of $\varepsilon_2$. When the initialisation is set to a previous solution, $\varepsilon_2 = 1 \times 10^{-6}$ is sufficient to find solutions that are visually indistinguishable from solutions with more restrictive tolerances.

*S3.3. Domain size*

Another important consideration for the numerical simulation is the choice of domain size, $L$. We aim to choose $L$ to be large enough that the agents do not touch the boundary during the simulations, but not so large as to incur a significant computational overhead.

To explore the choice of $L$ we always use the same number of nodes to discretise the nutrient equation, with $I^3 = 201^3$ equally–spaced nodes, and we set the first estimate of the solution for the GMRES algorithm to be the solution obtained by considering a different spheroid of the same radius but with a different placement of agents, similar to Supplementary S3.2, so that the computational efficiency of these tests is improved. We consider the domain lengths $L = 2000$, 4000, and $L = 6000\,\mu$m, and plot the resulting nutrient profiles along the midline $y = z = 0$ in Figure S6. While the boundary has a clear effect on the solution on the outer regions of the domain, the solutions are very similar in the region $|x| < 500\,\mu$m where agents are located and nutrient is being consumed. The two solutions for $L = 4000\,\mu$m and $L = 6000\,\mu$m are very similar in the region of the spheroid and also its immediate vicinity. Consequently, we choose a domain length $L = 4000\,\mu$m.

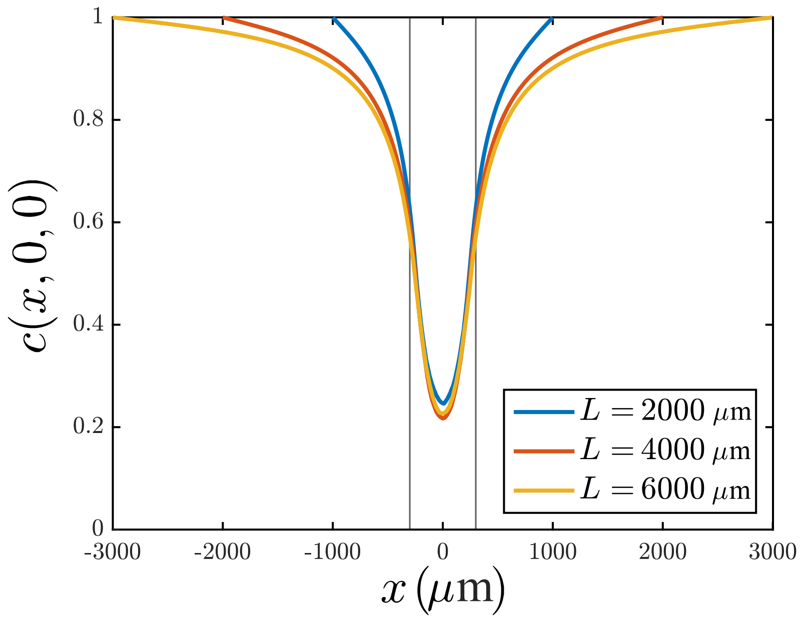

**Figure S6:** Comparison of steady-state solutions to Equation (S7) along one dimension ($x$) with varying domain lengths. The vertical lines represent the region of the test spheroid.

*S3.4. Spatial and temporal resolution of the nutrient profile*

There are two main considerations when we solve Equation (S7) for the nutrient profile: first we must choose an appropriate spatial discretisation, $I^3$; second, we must choose the duration of time between solutions of Equation (S7), $t^*$. Results in Figure S7 show the nutrient profiles along the midline $y = z = 0$ for a range of values of $I^3$ and $t^*$. The time scale for $t^*$ is hours, as this is the most appropriate choice for simulating agent-level behaviour, but we represent the results in days as a more intuitive and natural choice for growth on the scale of the spheroids.

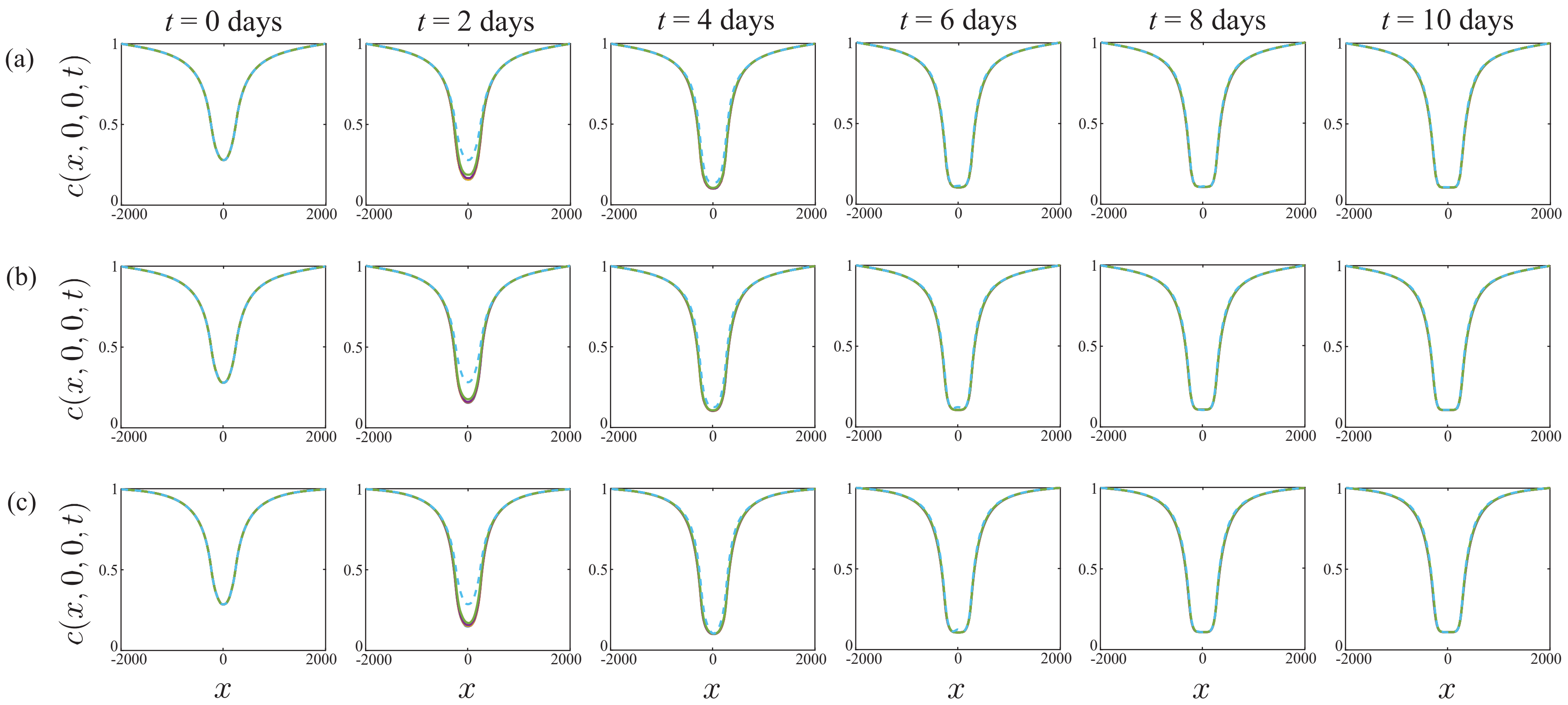

**Figure S7:** Influence of variation in $I^3$ and $t^*$ on nutrient solution over time. From left to right: nutrient profile solutions for day 0, 2, 4, 6, 8, and 10. Solution profiles show comparisons at these time points for $t^* = 1$ h (solid yellow), $t^* = 2$ h (solid purple), $t^* = 6$ h (solid green), and $t^* = 24$ h (dashed cyan). Solutions are averages of 10 simulations, shown for (a) $I^3 = 101^3$, (b) $I^3 = 151^3$, and (c) $I^3 = 201^3$.

Profiles in Figure S7 indicate that the solution is relatively insensitive to the choice of $I^3$, but we see a clear difference in solutions depending on the choice of $t^*$. The solutions in Figure S7 are insensitive to $t^*$ when we choose $t^* \leq 2$ h. Estimates of combinatorial runtime in Table S3 show that setting $t^* < 1$ h slows the simulations considerably. As a consequence, we choose $t^* = 1$ h.

**Table S3:** Table of average runtime data (min.) for 10 simulations with varying $I^3$ and $t^*$. The computational experimental period of these simulations is $T = 240$ h. Simulation runtimes are from high performance computing, using four 64 bit Intel Xeon core processors per simulation.

| $I^3$ | $t^* = 0.1$ h | $t^* = 0.5$ h | $t^* = 1$ h | $t^* = 2$ h | $t^* = 6$ h | $t^* = 24$ h |
|---|---|---|---|---|---|---|
| $101^3$ | 55.1 | 38.6 | 37.7 | 37.4 | 27.5 | 26.2 |
| $151^3$ | 101.0 | 57.1 | 50.8 | 48.4 | 37.6 | 36.3 |
| $201^3$ | 224.9 | 100.0 | 84.7 | 75.9 | 63.3 | 58.5 |

Additional results in Figure S8 show solutions for various grid resolutions $I^3 = 101^3$, $151^3$, $201^3$ for $t^* = 1$ h. At early time we see some difference in these solutions on the coarser mesh, but the solutions for $I^3 = 151^3$ and $201^3$ are visually indistinguishable at this scale. Therefore, in all simulations we set $t^* = 1$ h and $I^3 = 201^3$.

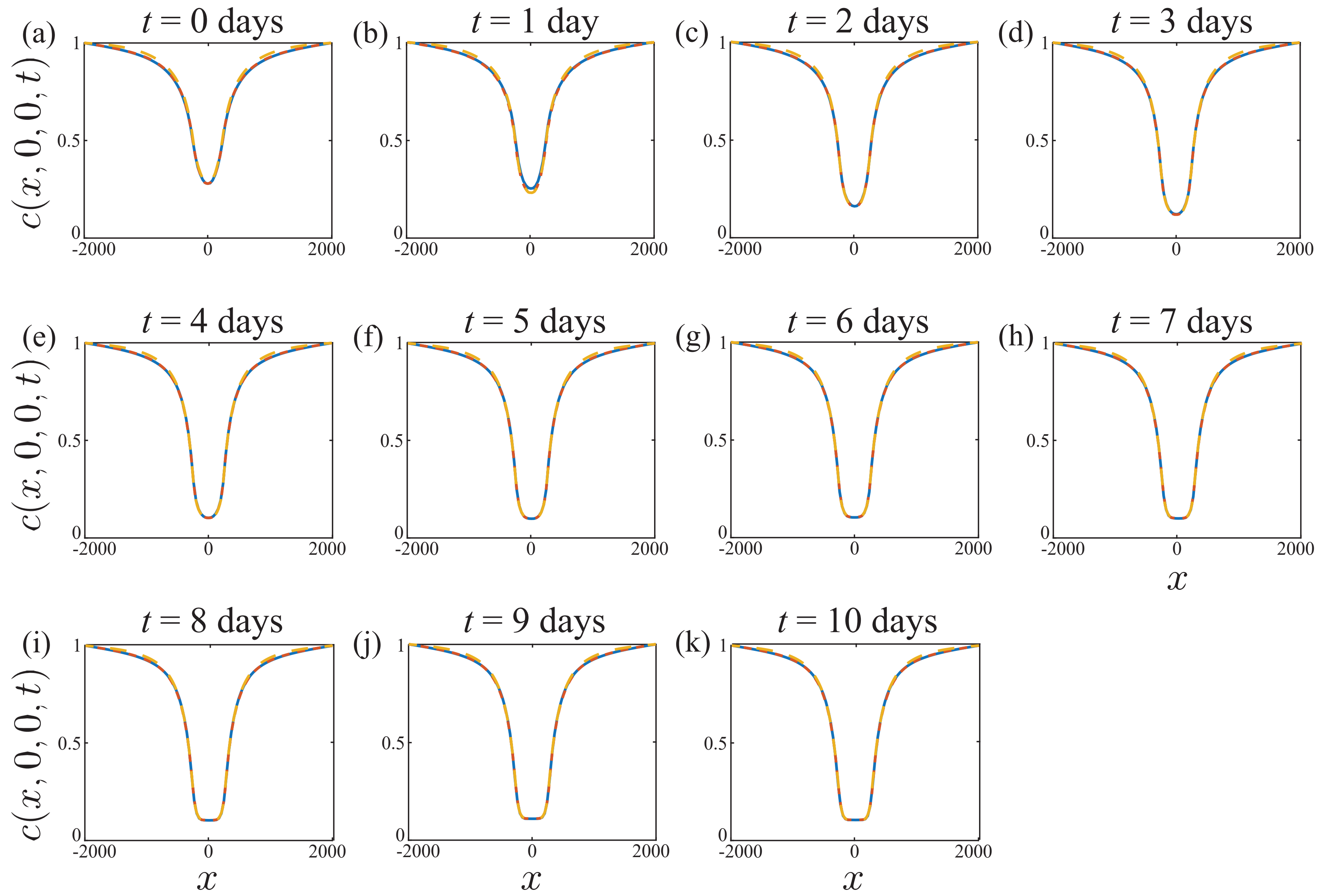

**Figure S8:** Influence of variation in $I^3$ when setting $t^* = 1$ h. (a)–(k) are comparisons of the solutions for $I^3 = 101^3$ (solid blue), $I^3 = 151^3$ (dashed red), and $I^3 = 201^3$ (dashed yellow) at days 0-10 respectively. As before, solutions are averages of profiles from 10 simulations.

### S4. Estimation of cell diameter

To reduce the number of unknown input parameters for the numerical experiments, we fix the experimental cell diameter $\Delta$, which in turn informs the proliferation and migration step size, $\sigma$ and $\mu$, respectively [7].

WM793B cells were stained with 10 $\mu$M CellTracker™ [8], at a density of $1 \times 10^6$ cells in 100 $\mu$L of tracker solution. After spinning the cell suspension at 300 relative centrifugal force for 2 minutes in a centrifuge, the CellTracker solution was removed and the pellet resuspended in 5 mL of growth medium. The cell suspension was spun down, before removal of the growth medium and resuspension of the stained cells in a solution of 2% low melting agarose in phosphate buffered solution (PBS). The solution was mounted on a chamber slide for preparation for imaging (Figure S9a). Images of the cell size were taken with a 20x Olympus UPlanSApo objective, where the Z range was set so that it was large enough to encompass a large sample of cells for analysis. Quantitative estimates of the WM793B cell size were achieved through computational image analysis with Fiji (ImageJ) software [9], using the 3D Cell Counter function to calculate the volume of the cells lying in the Z range.

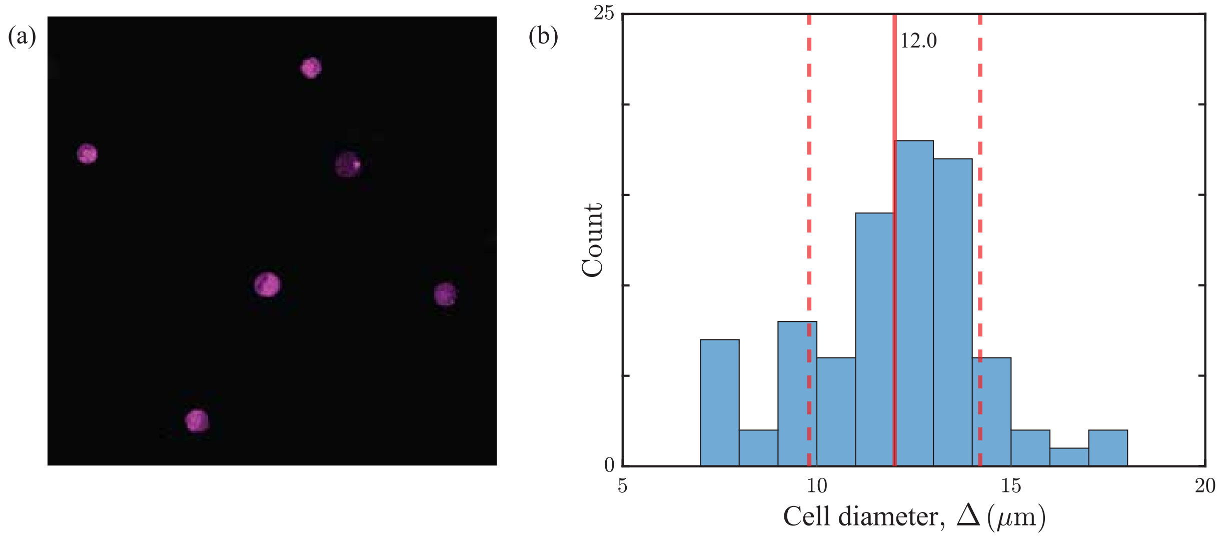

**Figure S9:** Estimation of the cell diameter for the WM793B cell line. (a) A visualisation of part of the window at one of the 50 Z heights, showing the equators of some of the suspended cells in the sample. The region identified with magenta colouring represents the entire cell. (b) Distribution of cell diameters from calculated volumes of the objects in the imaged volume. The solid line is the average diameter, and dashed lines represent one standard deviation from this average.

The 3D volume calculated by ImageJ is converted into an equivalent diameter of a sphere, giving $\Delta = 12.0\,\mu$m by taking the sample mean from 84 objects, distributed as shown in Figure S9b.

### S5. Estimation of cell cycle progression rates

We repeat the procedure for cell cycle duration estimates from [10] for the WM793b cell line. To estimate the maximum G1-early S phase (G1-eS) cycling rate $R_r$, the constant eS-S/G2/M cycling rate $R_y$, and the constant S/G2/M mitosis rate $R_g$, we use a series of 2D experiments as follows. The WM793B melanoma cells were seeded into a 12 well (3.82 cm$^2$ per well) #1.5 glass bottom plate, at a seeding density of 40 000 cells per well, and covered with 1 mL of growth medium, prepared as in [11]. The 2D cell culture was imaged in a Zeiss AxioObserver (Zeiss, Oberkochen, Germany) every 5 minutes for a period of 72 h, allowing for at least one full cell cycle to be completed. Then, 30 cells were tracked from the point of cell division through a full cell cycle, and the duration spent in each phase was measured and recorded. An example case of the fluorescence exhibited by individual cells in the two-dimensional assay is shown in Figure S10a.

Our measurements suggest that the average time spent in each stage of the cell cycle is $t_{\mathrm{r}} = 21.3$ h in red, $t_{\mathrm{y}} = 2.0$ h in yellow, and $t_{\mathrm{g}} = 16.2$ h in green. The progression rates are the reciprocals of these values, giving values of $R_r = 0.047\,/\mathrm{h}$, $R_y = 0.50\,/\mathrm{h}$, and $R_g = 0.062\,/\mathrm{h}$, which are the values for cell cycle progression that we use in the model. Figure S10b shows the time spent in the different phases of the cell cycle for each phase, with the mean and one standard deviation indicated.

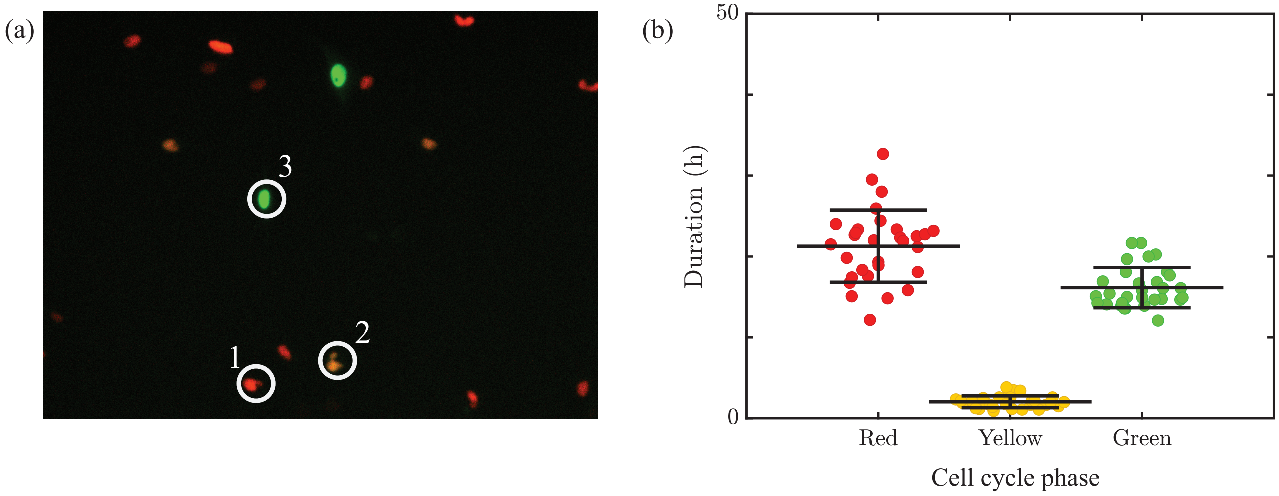

**Figure S10:** Estimation of the time spent in different stages of the cell cycle. (a) A section of one of the windows, indicating the visualisation and colourisation of the acquired FUCCI cell images. Examples of a red (1), yellow (2), and green (3) cell, where the yellow colour is generated by a composite of emission in the red and green channel. (b) A swarm chart of the duration of red, yellow, and green stages for 30 individual observations. Bars are mean ± standard deviation.

### S6. Initial cell number

Using the 3D experimental images, we estimate the initial cell count in the spheroid using FIJI (ImageJ) software [9] analysis of images at three different heights in the spheroid at $t = 0$ days. Once the image is loaded into ImageJ (Figure S11a), the image is binarised (Figure S11b) so that all pixels below a brightness threshold are black, and all above the threshold are white. Then, the watershed feature is used to segment connected groups of objects in the image, so that they may be counted properly (see Figure S11c-e). The finalised image is then processed with the Analyse Particles function, where the minimum size for an object is mandated to avoid counting residual pixels from the background removal. The minimum area for a cell was set at 25 $\mu$m$^2$. No maximum threshold was set, as experimentation with this parameter showed that it had a negligible effect on the cell count.

The area of the cells in the 2D image was recorded along with the cell count. Using the estimate of the cell diameter of 12 $\mu$m (from Supplementary S4), and assuming the density of cells in the slice area is the same as the volume density of cells in the entire spheroid, we obtain an approximate count of the cells in the spheroid at $t = 0$ days. This process was applied to images from the equator, and one from halfway to the top or bottom of the spheroid, labelled as the upper and lower cross sections (see Figure S11f). Two different spheroids at $t = 0$ days are used to estimate the total cell number at formation. Results (Table S4) indicate that spheroids at $t = 0$ days are composed of approximately 27 000 – 31 000 cells per spheroid. For simplicity, we simulate spheroids with 30 000 cells per spheroid at $t = 0$ days.

**Table S4:** Measurements of spheroid geometry and number of cells.

| Image name | Area ($\mu$m$^2$) | Radius ($\mu$m) | Volume ($\mu$m$^3$) | Cell count (area) | Density (%) | Cell count (spheroid) |
|---|---|---|---|---|---|---|
| Day 4(1) Equator | $1.58\times10^5$ | 224.24 | $4.72\times10^7$ | 739 | 52.91 | 27619 |
| Day 4(1) Upper cross section | $1.18\times10^5$ | 193.77 | $4.72\times10^7$ | 542 | 51.97 | 27128 |
| Day 4(1) Lower cross section | $1.14\times10^5$ | 190.15 | $4.72\times10^7$ | 511 | 50.88 | 26560 |
| Day 4(3) Equator | $1.84\times10^5$ | 242.24 | $5.95\times10^7$ | 766 | 46.99 | 30926 |
| Day 4(3) Upper cross section | $1.07\times10^5$ | 184.22 | $5.95\times10^7$ | 449 | 47.63 | 31343 |
| Day 4(3) Lower cross section | $1.58\times10^5$ | 223.94 | $5.95\times10^7$ | 643 | 46.16 | 30376 |

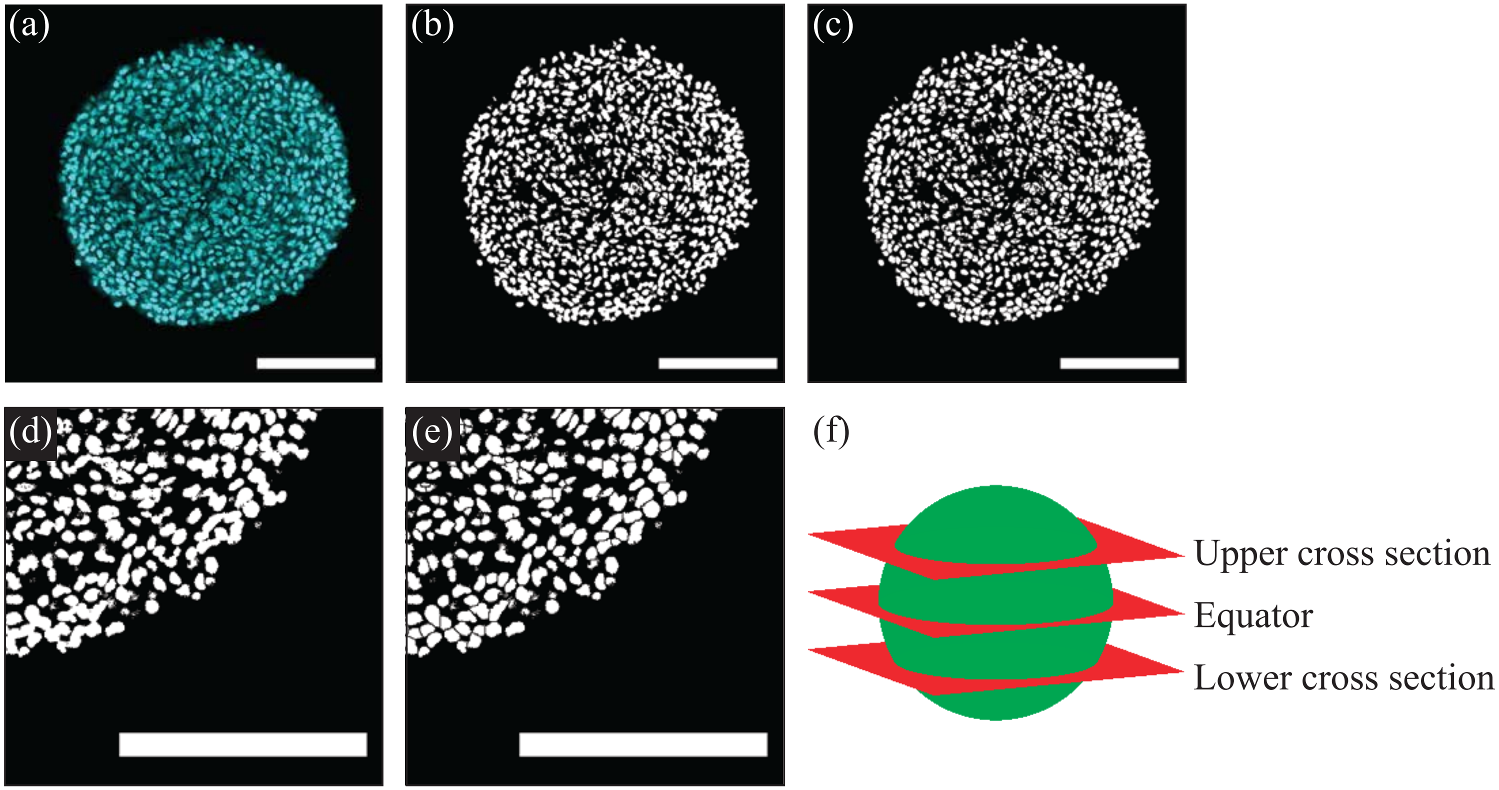

**Figure S11:** Demonstration of the process for preparing spheroid images for analysis. The spheroid is (a) imported into ImageJ, (b) binarised, and (c) watershed processed. (d)-(e) A zoomed-in demonstration showing the spheroid (d) before processing with watershed and (e) after processing with watershed. (f) Representation of the locations of the upper cross section, equator and lower cross section. Scale bars represent 200 $\mu$m.

### S7. Initialising the IBM

Here we describe how we choose the initial proportion of red, yellow, and green agents in each simulation. Results in Supplementary S5, show that the average time a cell spends in each phase of the cell cycle is $t_{\mathrm{r}} = 21.3$ h (red), $t_{\mathrm{y}} = 2.0$ h (yellow), and $t_{\mathrm{g}} = 16.2$ h (green). Therefore, we assume that the total cell cycle time is

$$t_{\mathrm{c}} = t_{\mathrm{r}} + t_{\mathrm{y}} + t_{\mathrm{g}}. \tag{S10}$$

We then define various population proportions $\beta_{\mathrm{r}} = t_{\mathrm{r}}/t_{\mathrm{c}}$, $\beta_{\mathrm{y}} = t_{\mathrm{y}}/t_{\mathrm{c}}$, and $\beta_{\mathrm{g}} = t_{\mathrm{g}}/t_{\mathrm{c}}$, for red, yellow, and green agents, respectively.

In the freely-cycling region of a spheroid, $r_{\mathrm{a}}(t) < r < r_{\mathrm{o}}(t)$, we distribute agent numbers according to these proportions. Since $\beta_{\mathrm{r}} + \beta_{\mathrm{y}} + \beta_{\mathrm{g}} = 1$, we can form a cumulative distribution of these agent proportions. For each agent, we sample a uniform random number $u_1 \in [0, 1]$, and perform inverse transform sampling on our cumulative distribution created by our population proportions, such that the agent is red if $u_1 < \beta_{\mathrm{r}}$, yellow if $u_1 \in [\beta_{\mathrm{r}}, \beta_{\mathrm{r}} + \beta_{\mathrm{y}}]$, and green otherwise (Figure S12a).

Experimentally, we see an initial arrested radius $r_{\mathrm{a}}(0) > 0$, meaning that the intensity of yellow and green for $r < r_{\mathrm{a}}(0)$ is less than 20% (Supplementary S1). Since we assume that G1-arrest is associated with low nutrient availability, we reduce the proportion of yellow and green agents to 16% of the population in the region where $c(\mathbf{x}, 0) < c_{\mathrm{a}}$, with the remaining 84% of the population specified to be red. Our choice of 16% is selected from a series of screening simulations that show this choice provides a good match to early experimental measurements. The algorithm implementing this approach is similar to that applied to agents in the freely-cycling region. For each agent with $c(\mathbf{x}, 0) < c_{\mathrm{a}}$ the agent is red with probability 0.84 (Figure S12b). Otherwise, we sample a uniform random number $u_2 \in [0, 1]$. Since we assume that commitment to the cell cycle (progression from G1 phase to eS phase) frees an agent from reliance on the local nutrient concentration, we expect that the yellow and green agents will be proportional to their cell cycle durations in this final 16% of the population, so that

$$\zeta_{\mathrm{y}} = \frac{t_{\mathrm{y}}}{t_{\mathrm{y}} + t_{\mathrm{g}}} \tag{S11}$$

is the proportion of the remaining 16% that is taken up by yellow agents, and

$$\zeta_{\mathrm{g}} = \frac{t_{\mathrm{g}}}{t_{\mathrm{y}} + t_{\mathrm{g}}} \tag{S12}$$

is the proportion of the 16% taken up by green agents. Hence, the agent is yellow if $u_2 < \zeta_{\mathrm{y}}$, and green otherwise (Figure S12b). Overall, we find that this experimentally-motivated approach to initialise our IBM simulations leads to a good match between simulated estimates of $r_{\mathrm{a}}(t)$ and our experimental measurements. Of course, if the IBM were applied to a model of 4D tumour spheroids with a different cell line, we anticipate that this approach would be valid, but that new measurements of $t_{\mathrm{r}}$, $t_{\mathrm{y}}$, and $t_{\mathrm{g}}$ would be required to apply this method.

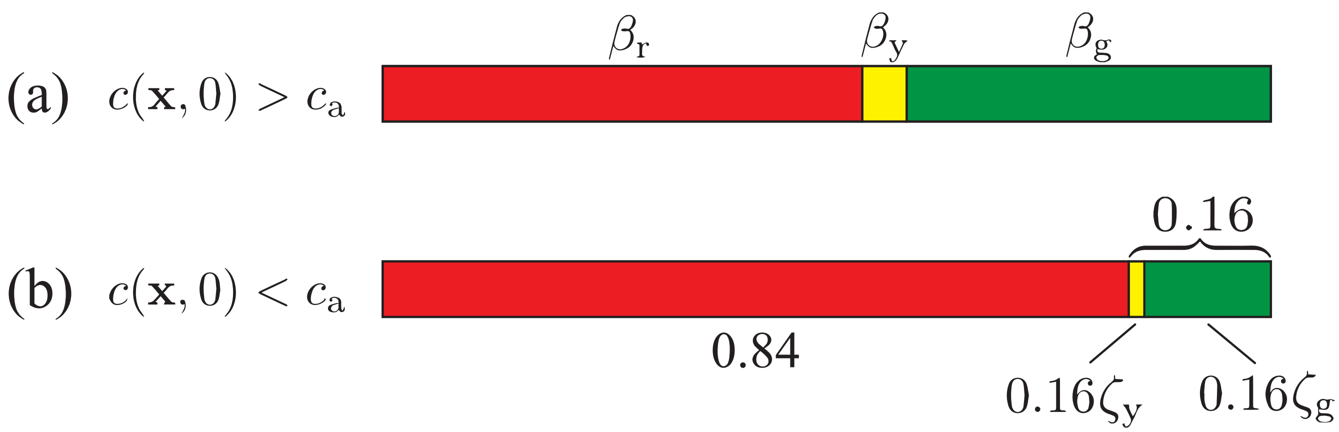

**Figure S12:** Representation of initial agent cell cycle stage proportions in (a) the freely-cycling region ($c(\mathbf{x}, 0) > c_{\mathrm{a}}$) and (b) the region of restricted nutrient concentration ($c(\mathbf{x}, 0) < c_{\mathrm{a}}$). In the freely-cycling region in (a), subpopulation proportions are set strictly by the proportions of time spent in each phase of the cell cycle, $\beta_{\mathrm{r}}$ for red agents, $\beta_{\mathrm{y}}$ for yellow agents, and $\beta_{\mathrm{g}}$ for green agents. In the region of restricted nutrient concentration in (b), the best qualitative match to experimental data is when red agents make up 84% of the population, and the remaining 16% is partitioned between yellow and green with proportions $\zeta_{\mathrm{y}}$ and $\zeta_{\mathrm{g}}$, respectively.

### S8. Simulation algorithms

Here, we describe pseudo-algorithms for the IBM, where Algorithm S1 solves the nutrient concentration with MATLAB code, and calls Algorithm S2 to calculate the agent-level behaviours with code in C.

**Algorithm S1:** A single realisation of the IBM, from $t = 0$ h to $t = T$ h.

Set parameter values from function inputs,
$\theta = (N(0), L, r, \sigma, \mu, T, t^*, R_r, R_y, R_g, d_{\text{max}}, d_{\text{min}}, m_{\text{max}}, m_{\text{min}}, \eta_1, \eta_2, \eta_3, I, \alpha, c_{\text{b}}, c_{\text{a}}, c_{\text{m}}, c_{\text{d}})$
Establish an $I^3$ equally-spaced finite volume mesh grid for the steady-state nutrient profile
Set $N(0)$ random initial agent locations within a sphere of radius $r$
Set $t = 0$
Solve the linear system of Equation (S7) for the initial agent density $\mathcal{C}$ with MATLAB's inbuilt GMRES function [5], setting `tol` $= 1 \times 10^{-8}$ ($\varepsilon_1$) and an initial guess of $c = 1$ at all nodes (Supplementary S3.2)
Calculate the nutrient concentrations from the mesh at the agent locations with linear interpolation
Assign each agent a cell cycle status (red, yellow, green), and record $N_{\text{r}}(0)$, $N_{\text{y}}(0)$, and $N_{\text{g}}(0)$ (Supplementary S7)
Calculate nutrient-dependent rates $R_r(c)$, $m(c)$, and $d(c)$ for all agents from Equation (**??**), Equation (**??**), and Equation (**??**)
**while** $t < T$
    Set $t = t + t^*$
    Solve the steady-state system of Equation (S7) with the current spheroid agent locations with MATLAB's inbuilt GMRES function, setting `tol` $= 1 \times 10^{-6}$ ($\varepsilon_2$) and an initial guess of $\boldsymbol{c}_{\text{p}}$, where $\boldsymbol{c}_{\text{p}}$ is the previous steady-state solution (Supplementary S3.2)
    Update $c_{\text{n}}$ for each agent with trilinear interpolation
    Resolve the cell events (Algorithm S2)
**end**

**Algorithm S2:** The Gillespie algorithm for simulating migration, cell cycle progression, and death.

Set inner timer $t_{\text{in}} = 0$

**while** $t_{\text{in}} < t^*$

$$d_{\text{t}} = \sum_{n=1}^{N(t)} (d(c))_n,\ m_{\text{t}} = \sum_{n=1}^{N(t)} (m(c))_n,\ R_{\text{t}} = \sum_{n=1}^{N_{\text{r}}(t)} (R_r(c))_n + N_{\text{y}}(t)R_y + N_{\text{g}}(t)R_g$$

Calculate the total rate of events $\lambda = d_{\text{t}} + m_{\text{t}} + R_{\text{t}}$

Sample the Gillespie time step to the next event $\tau \sim \text{Exp}(\lambda)$

Set $t_{\text{in}} = t_{\text{in}} + \tau$

Perform an event, with cycling probability $R_{\text{t}}/\lambda$, migration probability $m_{\text{tot}}/\lambda$, and death probability $d_{\text{tot}}/\lambda$

**if** *cycling event*

Sample agent $n$ to continue through the cell cycle, with probability proportional to its cycling rate

**if** *the agent is red*

Turn the agent yellow

$N_{\text{r}} = N_{\text{r}} - 1,\ N_{\text{y}} = N_{\text{y}} + 1$

Set the $n$th agent's cycling rate to $R_y$

**else if** *the agent is yellow*

Turn the agent green

$N_{\text{y}} = N_{\text{y}} - 1,\ N_{\text{g}} = N_{\text{g}} + 1$

Set the $n$th agent's cycling rate to $R_g$

**else if** *the agent is green*

Turn the agent red

Set $\theta = 2\pi u_1,\ \varphi = \arccos(1 - 2u_2);\ u_1,\ u_2 \sim \mathcal{U}(0, 1)$

$x_n = x_n + \sigma/2\cos(\theta)\sin(\varphi),\ y_n = y_n + \sigma/2\sin(\theta)\sin(\varphi),\ z_n = z_n + \sigma/2\cos(\varphi)$

$x_{\text{new}} = x_n - \sigma/2\cos(\theta)\sin(\varphi),\ y_{\text{new}} = y_n - \sigma/2\sin(\theta)\sin(\varphi),\ z_{\text{new}} = z_n - \sigma/2\cos(\varphi)$

$N_{\text{g}} = N_{\text{g}} - 1,\ N_{\text{r}} = N_{\text{r}} + 2$

Interpolate the nutrient concentrations to both new locations

Set $R_r(c)$, $m(c)$, and $d(c)$ for both new agents according to Equation (**??**), Equation (**??**), and Equation (**??**)

Adjust the population count in the agents' control volumes

**end**

**else if** *migration event*

Sample agent $n$ to migrate, with probability proportional to its migration rate

Set $\theta = 2\pi u_1,\ \varphi = \arccos(1 - 2u_2);\ u_1,\ u_2 \sim \mathcal{U}(0, 1)$

$x_n = x_n + \mu\cos(\theta)\sin(\varphi),\ y_n = y_n + \mu\sin(\theta)\sin(\varphi),\ z_n = z_n + \mu\cos(\varphi)$

Interpolate the nutrient concentration at the new location $\mathbf{x}_n$

Set $R_r(c)$, $m(c)$, and $d(c)$ for the agent at its new location with Equation (**??**), Equation (**??**), and Equation (**??**)

Account for changes to populations in control volumes, if necessary

**else if** *death event*

Sample an agent to die, with probability proportional to its death rate

$N = N - 1$

Remove the agent from the simulation, and move it to the dead population

**end**

**end**

### S9. Calculation of agent density profiles

Here, we describe the procedure to estimate the profiles of relative agent densities as functions of distance from the periphery in Figure **??**c. At a given time, $t$, we select the largest outer radius from all simulations, measured with image processing (Supplementary S1). We then calculate the largest multiple of 10 $\mu$m that is less than this selected radius, and denote this $r_{\mathrm{max}}$. For example, if at time $t$ the outer radii from all simulations are $r_{\mathrm{o}}(t)$ = 247, 248, 249, 251, and 252 $\mu$m, the largest outer radius is $r_{\mathrm{o}}(t) = 252\,\mu$m, and we choose $r_{\mathrm{max}} = 250\,\mu$m. This choice of $r_{\mathrm{max}}$ ensures that the bin corresponding to the largest radial distance does not include space beyond the spheroid periphery, and this is reasonable as, for 10 identically-prepared simulations, the variability in $r_{\mathrm{o}}(t)$ is less than 1% (Figure **??**).

We partition the interval $[0, r_{\mathrm{max}}]$ into bins of width $w = 10$ $\mu$m, which we find is sufficiently small to capture the spatial evolution of agent density, but is also sufficiently large to avoid excessive fluctuations [2]. The bins representing radial position are converted into bins representing distance from the periphery with the transformation,

$$\mathbf{p}_{\mathrm{b}} = \mathbf{r}_{\mathrm{max}} - \mathbf{r}_{\mathrm{b}}, \tag{S13}$$

where $\mathbf{r}_{\mathrm{max}}$ [$\mu$m] is a vector with all elements equal to $r_{\mathrm{max}}$ and length equal to the number of bin edges, and $\mathbf{r}_{\mathrm{b}}$ [$\mu$m] and $\mathbf{p}_{\mathrm{b}}$ [$\mu$m] are the locations of the bin edges for the radial distance and the distance to the periphery, respectively (Figure S13a).

We now demonstrate the calculation of the relative agent density distributions, using the green agents as an example, but the calculation is identical for the other agent types. First, all 3D Cartesian agent locations in a given simulation at $t$ are imported. We calculate the radial coordinate of the $n$th green agent, $r_n$, as the Euclidean distance from its 3D Cartesian coordinate, $\mathbf{x}_n$, to the mean location of all agents of all types at time $t$, $\bar{\mathbf{x}}(t)$, such that

$$r_n = ||\mathbf{x}_n - \bar{\mathbf{x}}(t)||, \quad n = 1, \ldots, N_{\mathrm{g}}(t), \tag{S14}$$

where $N_{\mathrm{g}}(t)$ is the number of green agents at time $t$. Assuming spherical symmetry, the distance to the spheroid periphery, $p_n(t)$, is calculated by subtracting $r_n$ from the outer radius, $r_{\mathrm{o}}(t)$,

$$p_n(t) = r_{\mathrm{o}}(t) - r_n, \qquad n = 1, \ldots, N_{\mathrm{g}}(t), \tag{S15}$$

where $p_n(t)$ is explicitly time-dependent through $r_{\mathrm{o}}(t)$. This calculation of the distance to the periphery for all green agents at time $t$ in a particular simulation is then repeated for all 10 simulations, and the sample means and standard deviations for these counts in each bin are calculated.

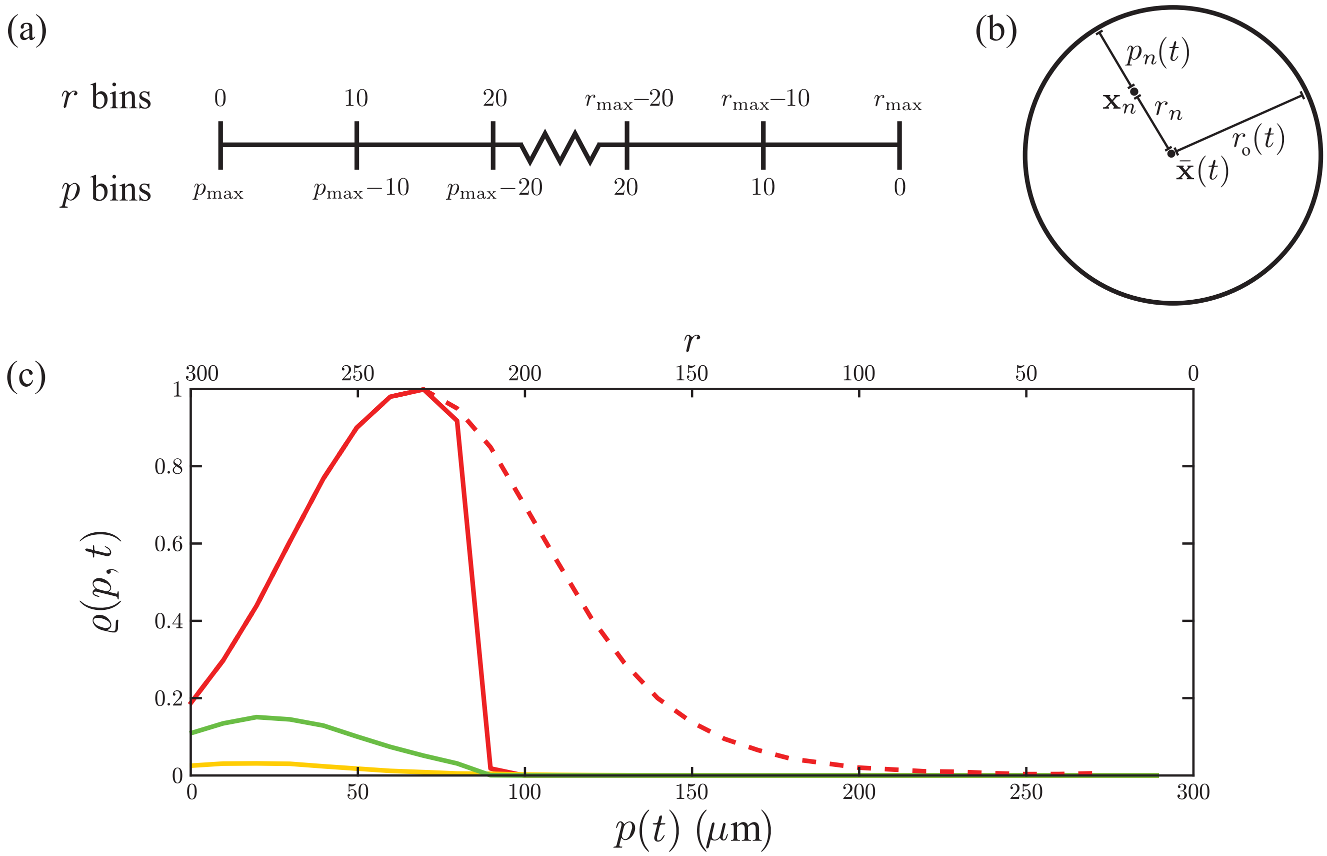

**Figure S13:** Schematic representation of the radial and distance to the periphery calculations. (a) The partitioning of bins for radial distance, and the associated bins for the distance to the periphery, calculated from the transformation in Equation (S13). (b) Demonstration of the positional calculations for radial coordinate and distance to the periphery under the assumption of spherical symmetry. An agent located at $\mathbf{x}_n$ within a spheroid at time $t$ with radius $r_{\mathrm{o}}(t)$ has its radial coordinate, $r_n$, calculated as the Euclidean distance to the average position of all agents, $\bar{\mathbf{x}}(t)$. This is then converted into the agent's distance from the periphery, $p_n(t)$, calculated from the transformation in Equation (S15) under the assumption of spherical symmetry. (c) An example plot of the relative agent density distributions, $\varrho(p, t)$, with the distance to the periphery on the lower axis, and the corresponding radial coordinate $r$ on the upper axis. We show example distributions of the cycling G1 (solid red), G1-arrested (dashed red), eS (yellow), and S/G2/M (green) relative agent densities.

We convert the average population counts by distance to the periphery, $C(p_l, t)$, into an agent density profile, $P(p_l, t)$ by dividing by the concentric spherical shell volume,

$$P(p_l, t) = \frac{3C(p_l, t)}{4\pi \left( \left( r_l + \frac{w}{2} \right)^3 - \left( r_l - \frac{w}{2} \right)^3 \right)}, \quad l = 0, \ldots, l_{\max}, \tag{S16}$$

where $p_l$ and $r_l$ are the centres of the $l$th bin for distance from the periphery and radius, respectively, and $l_{\max} = r_{\max}/w - 1$ is the number of bins. We then repeat the full procedure for the other agent types (cycling red, arrested red, yellow) to calculate the average agent density profiles for all agent types at $t$. We then repeat the calculation of $r_{\max}$, $\mathbf{p}_{\mathrm{b}}$, and density profiles for all agents at other values of $t$. All agent density profiles at all values of $t$ are then normalised with respect to a maximum density, $\varrho_{\max}$, so that, for each agent type,

$$\varrho(p_l, t) = \frac{P(p_l, t)}{P_{\max}}, \quad l = 0, \ldots, l_{\max}, \tag{S17}$$

where $P_{\max}$ is determined as the highest density of any type of agent over all values of $t$. This normalisation then gives the relative agent densities profiles, $\varrho(p, t)$ (Figure S13c), which are plotted in Figure **??**c at their corresponding time points.

### References

[1] Browning AP, Murphy RJ, 2021. Image processing algorithm to identify structure of tumour spheroids with cell cycle labelling. *Zenodo*. doi: 10.5281/zenodo.5121093.

[2] Binder BJ, Simpson MJ, 2015. Spectral analysis of pair-correlation bandwidth: application to cell biology images. *Royal Society Open Science*, **2**:140494. doi: 10.1098/rsos.140494.

[3] Crank J, 1975. *The Mathematics of Diffusion*. Clarendon Press, Oxford, 2nd edition.

[4] Mathworks, 2021. MATLAB `ode45`. https://www.mathworks.com/help/matlab/ref/ode45.html. (Accessed: November 2021).

[5] Mathworks, 2021. MATLAB `gmres`. https://www.mathworks.com/help/matlab/ref/gmres.html. (Accessed: November 2021).

[6] Mathworks, 2021. MATLAB `mldivide`. https://au.mathworks.com/help/matlab/ref/mldivide.html. (Accessed: November 2021).

[7] Simpson MJ, Landman KA, Hughes BD, 2010. Cell invasion with proliferation mechanisms motivated by time-lapse data. *Physica A: Statistical Mechanics and its Applications*, **389**:3779–3790. doi: 10.1016/j.physa.2010.05.020.

[8] ThermoFisher Scientific, 2021. Celltracker™. https://www.thermofisher.com/order/catalog/product/C34565. (Accessed: November 2021).

[9] Schindelin J, Arganda-Carreras I, Frise E, Kavnig V, Longair M, Pietzsch T, Preibisch S, Rueden C, Saalfeld S, Schmid B, et al., 2012. Fiji: an open-source platform for biological-image analysis. *Nature Methods*, **9**:676–682. doi: 10.1038/nmeth.2019.

[10] Haass NK, Beaumont KA, Hill DS, Anfosso A, Mrass P, Munoz MA, Kinjyo I, Weninger W, 2014. Real-time cell cycle imaging during melanoma growth, invasion, and drug response. *Pigment Cell & Melanoma Research*, **27**:764–776. doi: 10.1111/pcmr.12274.

[11] Spoerri L, Beaumont KA, Anfosso A, Haass NK, 2017. Real-time cell cycle imaging in a 3D cell culture model of melanoma. *Methods in Molecular Biology*, **1612**:401–416. doi: 10.1007/978-1-4939-7021-6_29.
